# Maximum likelihood pandemic-scale phylogenetics

**DOI:** 10.1101/2022.03.22.485312

**Authors:** Nicola De Maio, Prabhav Kalaghatgi, Yatish Turakhia, Russell Corbett-Detig, Bui Quang Minh, Nick Goldman

## Abstract

Phylogenetics plays a crucial role in the interpretation of genomic data^1^. Phylogenetic analyses of SARS-CoV-2 genomes have allowed the detailed study of the virus’s origins^2^, of its international^3,4^ and local^4–9^ spread, and of the emergence^10^ and reproductive success^11^ of new variants, among many applications. These analyses have been enabled by the unparalleled volumes of genome sequence data generated and employed to study and help contain the pandemic^12^. However, preferred model-based phylogenetic approaches including maximum likelihood and Bayesian methods, mostly based on Felsenstein’s ‘pruning’ algorithm^13,14^, cannot scale to the size of the datasets from the current pandemic^4,15^, hampering our understanding of the virus’s evolution and transmission^16^. We present new approaches, based on reworking Felsenstein’s algorithm, for likelihood-based phylogenetic analysis of epidemiological genomic datasets at unprecedented scales. We exploit near-certainty regarding ancestral genomes, and the similarities between closely related and densely sampled genomes, to greatly reduce computational demands for memory and time. Combined with new methods for searching amongst candidate evolutionary trees, this results in our MAPLE (‘MAximum Parsimonious Likelihood Estimation’) software giving better results than popular approaches such as FastTree 2^17^, IQ-TREE 2^18^, RAxML-NG^19^ and UShER^15^. Our approach therefore allows complex and accurate probabilistic phylogenetic analyses of millions of microbial genomes, extending the reach of genomic epidemiology. Future epidemiological datasets are likely to be even larger than those currently associated with COVID-19, and other disciplines such as metagenomics and biodiversity science are also generating huge numbers of genome sequences^20–22^. Our methods will permit continued use of preferred likelihood-based phylogenetic analyses.

## Main

As viruses and bacteria spread within and between hosts, they accumulate genetic mutations. By analysing the genetic data of sampled pathogens, we can understand their evolutionary and transmission history. For this reason, genomic data play a crucial role in epidemiology, as exemplified during the COVID-19 pandemic, and are used to track and reconstruct the spread of disease within communities and within and between countries^4–6,23–25,^ understand the dynamics of transmission^7,24,26,27^, estimate the efficacy of containment measures^3,28–30^, predict future epidemiological dynamics^9,23^, and for the tracking of pathogen evolution as showcased by the identification of new SARS-CoV-2 mutations and variants of concern^8,10,11,31,32^.

Investigations of genomic epidemiological data are predominantly based on phylogenetic methods, but analyses of SARS-CoV-2 genome sequence data with existing phylogenetic approaches are becoming more difficult due to the excessive computational resources required by current global datasets consisting of millions of genomes^16^. While a daily updated global SARS-CoV-2 phylogenetic tree is particularly useful^33^, estimating it with established phylogenetic software like RAxML^34^ or IQ-TREE^18^ would require years for each tree update (if possible at all due to memory demand). For this reason, tools for tracking viral genome evolution and spread (e.g. NextStrain^35^) and many other genomic analyses often downsample global SARS-CoV-2 datasets to a few thousand genomes, leading to loss of power and resolution^36,37^.

### A new approach for pandemic-scale likelihood-based phylogenetics

To address these issues, we have devised a set of new algorithms, techniques and formats tailored for large-scale genomic epidemiology. Our approach, MAPLE (“MAximum Parsimonious Likelihood Estimation”), performs maximum likelihood (“ML”) phylogenetic inference^17,18,34^ and uses explicit probabilistic models of sequence evolution; we combine these best-in-class features with some aspects of maximum parsimony methods^15^ that allow it to greatly reduce computer memory and time demand.

#### Concise genome data representation

Genomic data typically need to be aligned before performing phylogenetic inference; resulting alignments usually employ Fasta or similar formats^38^ which list the whole DNA sequence of each considered sample. In the context of genomic epidemiology this is very highly redundant since genomes within an epidemic are usually extremely similar to each other. While it is possible to reduce the size of datasets using standard compression techniques^39^, sequences still need to be uncompressed before analysis.

Instead, we represent each genome in our MAPLE alignment format in terms of differences with respect to a reference genome (Fig. 1A and Methods). This way, we reduce file size approximately 100-fold compared to Fasta files (Extended Data Fig. S1); for example, we reduced the size of the 31-03-2021 GISAID global SARS-COV-2 alignment of 915,508 genomes from 27.84 GB to 224.6 MB (a 124× reduction).

**Figure 1:**
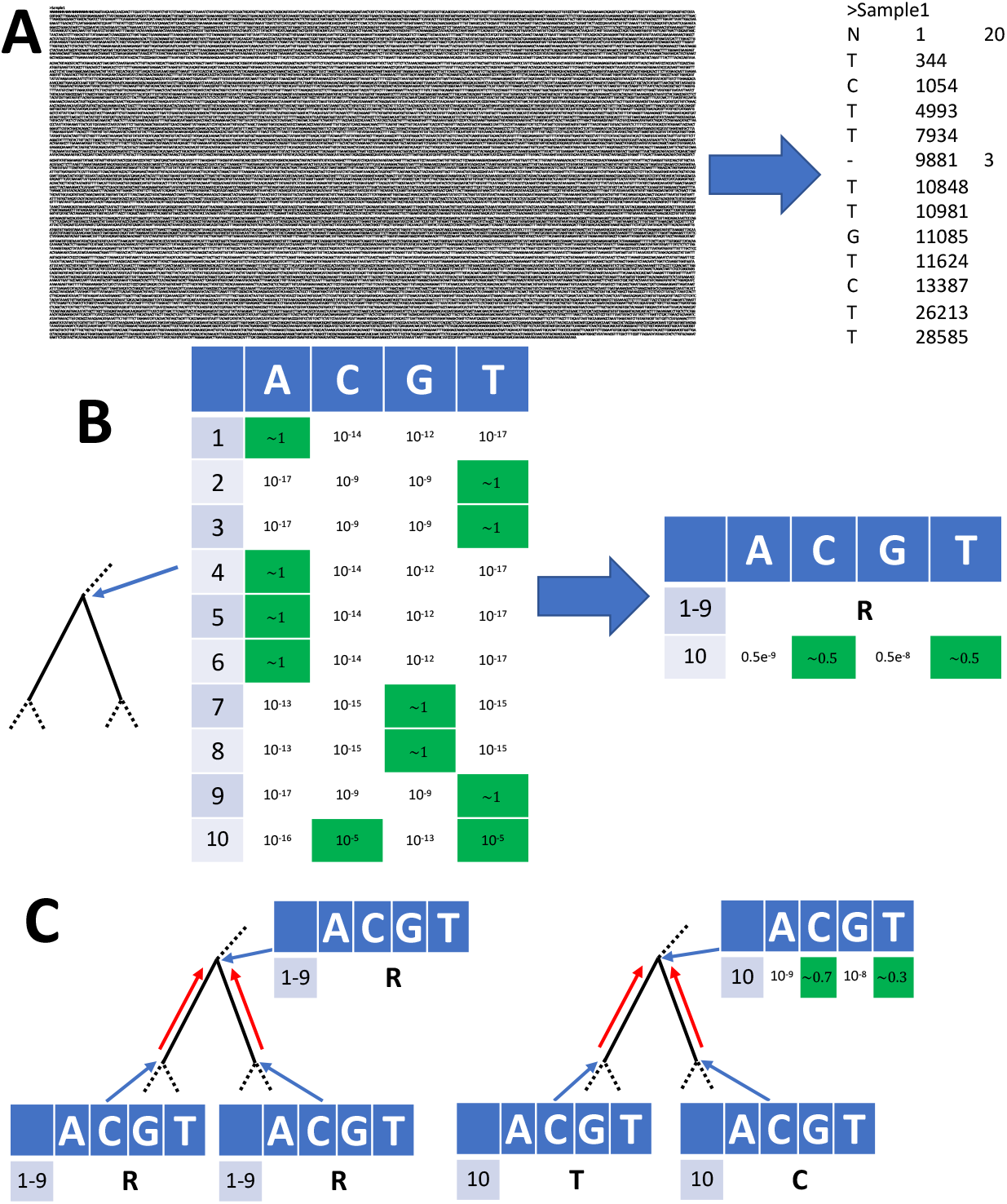
Graphical summary of sequence and likelihood representation and processing. **A** Left: Fasta representation of an individual SARS-CoV-2 genome consists of sample name followed by the entire ≈30 kbp genome sequence. Right: MAPLE format records only the differences between the genome under consideration and a reference; columns represent the variant character observed, the position along the genome, and (when necessary) the number of consecutive positions for which the character is observed. **B** Left: an example likelihood vector at an internal node of a phylogenetic tree (shown by the narrow blue arrow; only a small portion of the tree is shown); for simplicity we show only 10 genome positions. At each position (rows 1–10), each column contains the likelihood for a specific nucleotide. For rows 1–9 the likelihood is concentrated at only one nucleotide (highlighted in green), while for position 10 we show an example with more uncertainty. Right: MAPLE representation of these node likelihoods. Assuming that the reference sequence at the first 9 positions matches the most likely nucleotides in the vector (ATTAAAGGT) then for positions 1–9 the likelihood of non-reference nucleotides is negligible and we represent the likelihoods with a single symbol (R). At position 10, due to non-negligible uncertainty, we explicitly calculate and store the four relative likelihoods. **C**: Examples of likelihood calculation steps in MAPLE. Red arrows represent the flow of information from the tips to the root of the tree. Left: if two child nodes are in reference state R for a region of the genome (here, positions 1–9), then MAPLE assumes that their parent is also in state R. Right: if at a genome position (here, position 10) two child nodes have likelihoods concentrated at different nucleotides, then for their parent we explicitly calculate the relative likelihoods of all four nucleotides.

#### Concise phylogenetic likelihood representation and calculation

Likelihood-based phylogenetic methods typically keep track of the probability of every possible nucleotide at each position of the genome and each node of the phylogenetic tree^1,14^. With pandemic-scale genomic data, this process requires excessive computational time and memory resources^16^. However, in genomic epidemiology, due to the similarity of the genomes considered, these probabilities are typically highly concentrated at only one of the four nucleotides for most genome positions and tree nodes. We exploit this feature by approximating these probabilities and representing them concisely (Fig. 1B and Methods). As an example, when estimating a phylogeny from a random 10,000-sample subset of the GISAID dataset above, with a reference genome of 29,891 bp, on average we only record the phylogenetic likelihoods of 2.7 genome positions per tree node (≈10,000 times less than usual). This allows us to considerably reduce the memory demand of likelihood-based phylogenetic inference in genomic epidemiology.

Additionally, we develop a more efficient alternative to the Felsenstein pruning algorithm^14^ used to calculate phylogenetic likelihoods; this algorithm has been at the core of most of likelihood-based phylogenetics in the past 40 years, and so is fundamental to some of the most cited and used scientific software, but is not tailored for the features of pandemic-scale genomic data. Our alternative (Fig. 1C and Methods) takes advantage of the strong similarities between the considered genomes and of efficient likelihood and data representation to reduce the computational time demand of likelihood-based phylogenetics in genomic epidemiology.

#### Efficient tree exploration

To efficiently but accurately find likely phylogenetic trees, we develop new strategies for exploring tree space. Our first strategy is an adaptation of stepwise addition^40^, in which samples are added to the phylogenetic tree one at the time. We use this strategy to find an initial tree (which is then refined with the second strategy), but it is similarly useful in extending an existing tree, for example as new genomes become available with time. Our adaptation involves an efficient search among the nodes of the tree for the most likely tree position in which to add the new sample (Fig. 2 and Methods).

**Figure 2:**
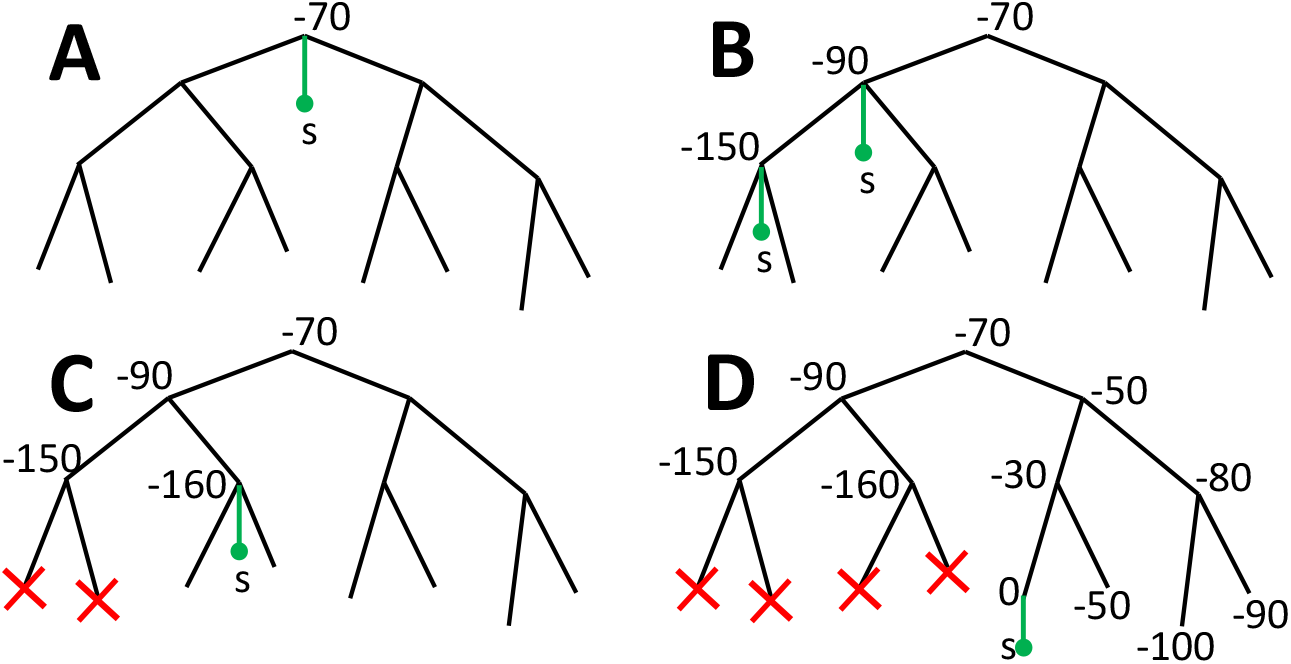
Graphical summary of phylogenetic placement in MAPLE. **A** To search for the best placement of a new sample s (here represented by a green dot and branch) on the current tree, we first assess placement at the root, which in this case results in a relative log-likelihood score of -70. **B** We iteratively visit descendant nodes by preorder traversal and assess placement for each visited node (in practice we also attempt placement onto branches). **C** When the log-likelihood score decreases two times consecutively and falls below a certain threshold relative to the best placement found so far, we do not visit further nodes downstream (red crosses). **D** The placement with the highest score at the end of this process (in this case with cost 0) is taken as optimal for the addition of s to the tree.

Our second strategy consists of a modification of subtree pruning and regrafting^40^, which is used to perturb (and thereby improve) an existing tree. Our modification consists again in efficiently exploring a broad range of possible tree changes.

### Computational demand and accuracy of MAPLE

Maximum likelihood phylogenetic methods typically present trade-offs between accuracy and computational demand, with more accurate tree reconstruction requiring deeper, and therefore more time consuming, tree space exploration. Thanks to the considerable time and memory savings brought by our new approach to likelihood calculation, MAPLE can invest more resources in tree estimation than other methods, resulting in more accurate tree inference, while requiring less time and memory than other maximum likelihood inference approaches (Fig. 3, Extended Data Figs. S2–S5).

**Figure 3:**
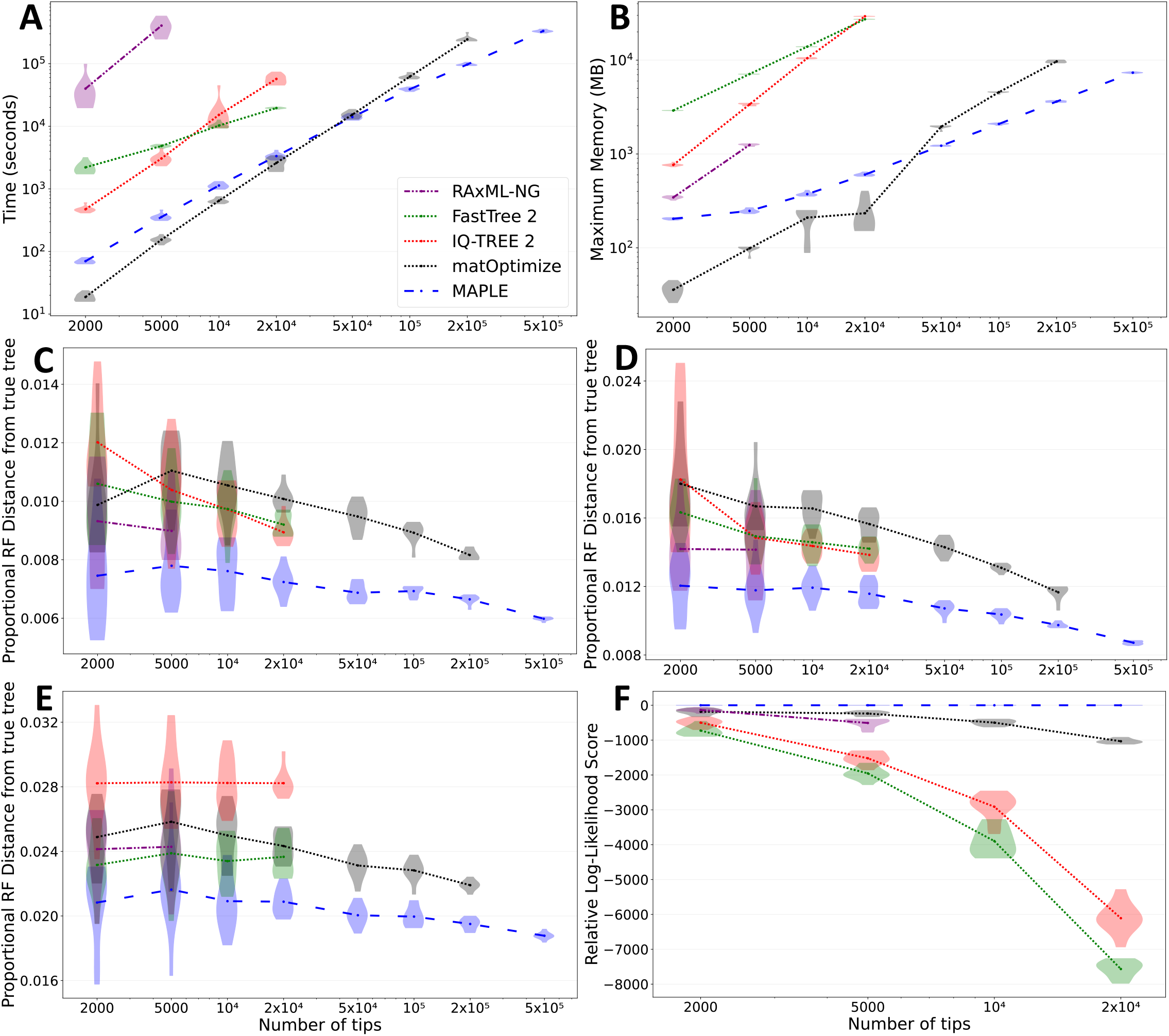
MAPLE consistently delivers higher accuracy phylogenetic inference from SARS-CoV-2 genomes at lower computational demand. **A** Time required to run each method considered on real SARS-CoV-2 datasets. Each phylogenetic inference method considered is represented by a different color and line style (see legend). Values on the X axes show the number of samples included in each replicate. We ran each method up to the maximum dataset size that could be analysed due to time (one week) and memory (40GB) limitations. Each violin plot summarizes values for 10 replicates, and dots represent mean values. **B** Maximum RAM demand required to run each method considered on real SARS-CoV-2 datasets. **C–E** Proportional Robinson-Foulds distances between estimated trees and true trees in simulations. Higher values correspond to more errors in phylogenetic estimation. **C** “Basic” simulation scenario; **D** “rate variation” simulation scenario; **E** “sequence ambiguity” simulation scenario. **F** Log-likelihoods of phylogenies inferred by different methods on real SARS-CoV-2 data, relative to the highest log-likelihood score obtained by any method for the same replicate. Higher values on the Y axis represent more likely estimates. We consider only datasets of up to 20,000 samples due to the computational demand of likelihood evaluation.

As an example, MAPLE shows consistently higher accuracy than RAxML-NG^19^ (the most accurate of the methods we compared MAPLE against) on simulated and real SARS-CoV-2 datasets (Fig. 3C-F, Extended Data Figs. S4 and S5), while being more than 100-fold faster (Fig. 3A) and requiring less memory (Fig. 3B). MAPLE can also estimate trees about 25 times larger than IQ-TREE 2^18^ or FastTree 2^17^ (500,000 vs 20,000 samples) because of their 50-fold larger memory demand (Fig. 3B). Fig. 4 shows an example 500,000-sample SARS-CoV-2 whole-genome phylogeny inferred by MAPLE.

**Figure 4:**
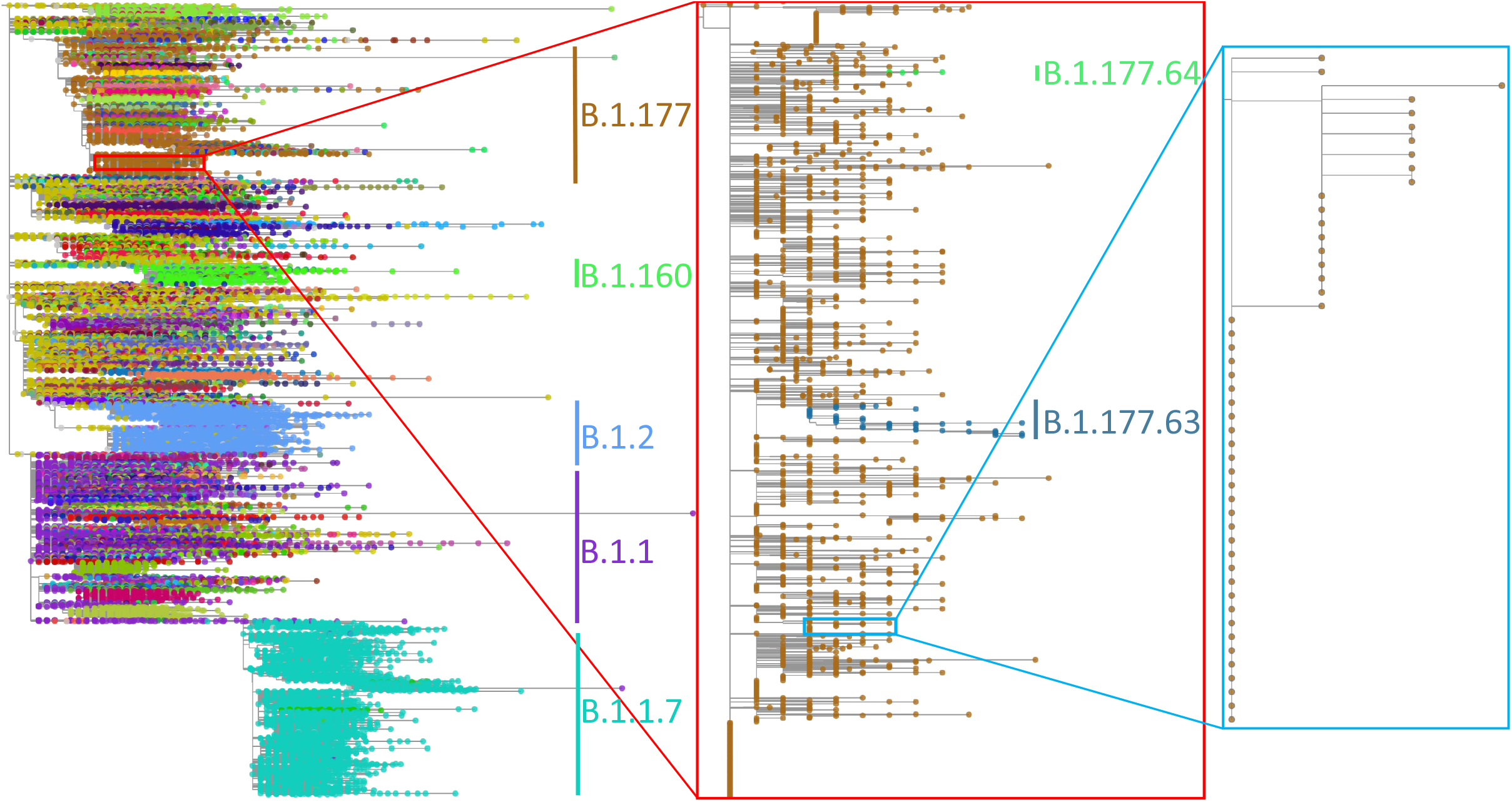
500,000-sample phylogeny inferred by MAPLE. Example phylogeny, with two consecutive zoom-ins each of about 100x magnification. Different SARS-CoV-2 lineages are shown in different colors, with some clades labelled to give context. Left: 500,000-sample phylogeny estimated by MAPLE from real SARS-CoV-2 sequence data. Center: Zoom-in on a subtree containing 3,600 B.1.177 samples. Right: Further zoom-in on a subtree containing 49 samples. Phylogenies plotted using Taxonium^42^.

matOptimize^41^ (a recent feature improving the accuracy of UShER^15^) is a phylogenetic inference method that, similarly to MAPLE, has been tailored to the features of genomic epidemiological analyses, but that uses maximum parsimony rather than maximum likelihood principles. MAPLE shows similar computational demand to matOptimize, and less steep slopes in time and memory demand, being therefore able to estimate larger trees (Fig. 3A-B). matOptimize appears less accurate than maximum likelihood methods on simulated data (Fig. 3C-E), but more accurate on real data (Fig. 3F), being second only to MAPLE.

We can further improve the computational performance of MAPLE by reducing the depth of its tree space search; for example, using option “- -fast” in MAPLE, runtime becomes typically 2-3 times faster (Extended Data Fig. S2) without decreasing accuracy on simulated datasets (Extended Data Fig. S4) and while remaining the most accurate approach on real data (Extended Data Fig. S5).

## Discussion

By rewriting the classic Felsenstein pruning algorithm, by including features of parsimony-based phylogenetic inference in a likelihood-based context, by using efficient approximations, and by using more concise data representation, we have achieved substantial reductions in memory and time demand and increases in accuracy compared to popular ML approaches when inferring SARS-CoV-2 phylogenies. This enables state-of-the-art phylogenetic inference to be performed on larger datasets than previously possible.

Beyond SARS-CoV-2, our approach will be equally useful in any analysis with many sequences and with short evolutionary distances, such as most scenarios in genomic epidemiology. This includes genomic datasets with many samples from an individual pathogen, including for example large collections of *M. tuberculosis* genomes^43^ or influenza genomes^44^, and collections of genomic data from possible future pandemics. Our approach could also be combined with divide-and-conquer phylogenetic algorithms^45,46^ to further improve its performance and applicability.

The applicability of our methods goes beyond ML phylogenetics. The same algorithms and data structures in MAPLE could also be used in a Bayesian setting, since Bayesian phylogenetic methods (for example BEAST^47,48^) use the same genetic data (multiple sequence alignments) and the same likelihood calculation algorithms as ML phylogenetic methods, and so would benefit from the same reduction in computational demands.

For these reasons, we expect that in the future MAPLE and its algorithms will expand the computational toolkit of genomic epidemiology and could improve our preparedness for combating future epidemics.

## Methods

### Concise representation of genomic epidemiological sequence data

We use a concise and human-readable format for representing an alignment of closely related genome sequences, which we call MAPLE format. We express each genome sequence in terms of its differences (substitutions and deletions) with respect to the reference. We also record ambiguous positions (IUPAC ambiguity characters), and deleted or non-sequenced portions of the genomes (gap “-” and “N” characters, respectively).

As an illustrative example, we consider a reference genome “Reference” comprising 20 “A” characters:

~~~
>Reference
AAAAAAAAAAAAAAAAAAAA
~~~

(here represented in Fasta format). If a sampled genome “Sample” consists of the sequence:

~~~
>Sample
NNNNNAAAAA---AAAAATA
~~~

when aligned to the reference, as it would be represented in Fasta format, we instead represent it as:

~~~
>Sample
N 1 5
- 11 3
T 19
~~~

where in each entry (row) the first column represents the type of difference with respect to the reference, the second column in each row represents the position (along the reference genome) of the difference, and the third column (which we only require for “N” and “-” entries) represents how many consecutive positions have this same character.

### Concise representation of ancestral sequences and sequence uncertainty

In addition to representing sequence data efficiently, we also efficiently calculate and represent partial likelihoods at internal nodes of the tree. Given a node *n* of the phylogenetic tree *ϕ*, a column *i* of alignment *A* containing site pattern (nucleotides) *A*_*i*_, and an evolutionary model *M*, the partial likelihood at *n* and *i* of nucleotide *X* is typically defined in phylogenetics as

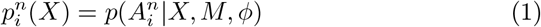

where 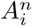 is the subset of observations in *A*_*i*_ corresponding to the descendant leaf nodes of *n*. These partial likelihoods are typically calculated with the Felsenstein pruning algorithm^1^; in total, there are 4 ×*L*× |*ϕ*| such likelihoods that need to be computed, stored and repeatedly updated during phylogenetic inference, where *L* is genome length and |*ϕ*| is the number of nodes in *ϕ*. For SARS-CoV-2, *L* >29,000 bp and |*ϕ*| can be in the order of millions, making this approach unfeasible.

Instead, we replace partial likelihood vectors with more concise structures that we call “genome lists”. Each entry of a genome list represents phylogenetic partial likelihoods for either one position of the genome or for a set of consecutive positions that share similar features. An important difference from the traditional Felsenstein pruning method is that, for each genome position and tree node, we only keep track of relative partial likelihoods among the four nucleotides, and not exactly of each 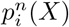; in other words, we aim at tracking values 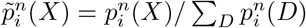. An entry of our genome list is a tuple of four elements (*τ, i, l, v*), comprising:

- an entry “type” *τ*; the permitted types are “**R**”, to indicate collections of contiguous sites that are identical to the reference (that is, sites where the partial likelihoods are all concentrated at the reference nucleotide); type “**N**” to indicate contiguous sites that contain no descendant sequence information (that is, sites where all four nucleotides have the same partial likelihoods); types “**A**”, “**C**”, “**G**” and “**T**” to indicate individual sites where the corresponding non-reference nucleotide is the ancestral one at the node with negligible uncertainty (that is, the partial likelihood mass is all concentrated in one non-reference nucleotide); and type “**O**” (“other”) to indicate positions where multiple nucleotides have non-negligible relative partial likelihoods
- a “position” *i* representing the position of the reference to which the entry refers. If the entry corresponds to a stretch of sites, this element is the position of the first one (from 5′ to 3′) of these. The last position of the entry need not be specified explicitly
- the “branch length” *l* represents the evolutionary distance (using the same unit used to represent branch lengths) between node *n* and the location in the tree where the partial likelihoods contained or represented by the genome list entry refer to (see e.g. Extended Data Fig.S6C). *l* efficiently carries information regarding the uncertainty of sites’ states by recording the evolutionary distance from the last visited position in the tree with no state uncertainty.
- relative partial likelihoods (“partials”) *v*, representing the vector 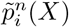 for the position considered — only needed for entries of type “**O**”.

Where we have made use of the concept of negligibility to distinguish entries of type “**O**” from the others, in practice we define negligibility through an arbitrary threshold *ϵ* with default value *ϵ* = 10^*-*8^, that is, a site is of type “**O**” only if at least two nucleotides have a relative partial likelihood 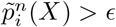.

As an example, we can consider the sample in the previous section

~~~
>Sample
N 1 5
- 11 3
T 19
~~~

and the same reference genome comprising 20 “A” nucleotides. Under these assumptions, at the terminal node of the phylogeny corresponding to “Sample”, we have the genome list

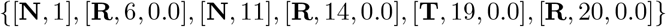

We omit branch length elements (third elements in each entry) of entries of type “**N**” since they are redundant.

If instead of a “T” character at position 19 we observed a IUPAC ambiguity code^2^ “Y” (meaning “C” or “T”), then the fifth entry of the genome list would have been

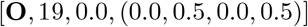

### Calculation of genome lists

As is standard in phylogenetics, we assume that sequence evolution is a continuous-time and finite-space homogeneous Markov process, where all sites evolve independently^3^. We assume a nucleotide substitution process determined by a substitution rate matrix *Q* whose entries *q*_*XY*_, for any *X* ≠ *Y*, represent instantaneous rates of substitution of nucleotide *X* to nucleotide *Y*, and *q*_*XX*_ = − Σ_*Y ≠ X*_ *qXY*. Transition probabilities over a branch length *l* are typically calculated using matrix exponentiation^3^, for which, considering the short branch lengths involved in genomic epidemiology, we use a first order approximation:

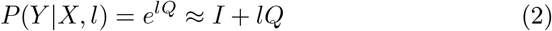

where *I* is the identity matrix. This means that the probability *P* (*Y* | *X, l*) of nucleotide *X* evolving into nucleotide *Y* ≠ *X* is approximated as *lq*_*XY*_, and that *P* (*X* | *X, l*) ≈ 1+ *lq*_*XX*_.

For simplicity, we assume that the tree *ϕ* is binary and rooted, that is, each internal node has exactly two children. We represent multifurcations using bifurcations separated by branches of length 0.

Similarly to the Felsenstein pruning algorithm, we calculate the genome list of an internal node *n* only after calculating it for its children. We have shown above how we initialize genome lists for terminal nodes of the tree. Now, we assume that *n* has child nodes *b*_1_ and *b*_2_ with genome lists respectively *L*_1_ and *L*_2_. We also assume that *b*_1_ and *b*_2_ are separated from *n* by branches of length *l*_1_ and *l*_2_. We want to calculate the genome list *L*_n_ of node *n*, which we obtain by “merging” information from *L*_1_ and *L*_2_.

Given the two genome lists *L*_1_ and *L*_2_, we split the genome into segments, where each segment corresponds to genome positions that all belong to the same genome list entry in *L*_1_, and also all belong to the same entry in *L*_2_ (see Supplementary Methods Section S1.1 for more details). For example, if we assume our usual reference of 20 “A” nucleotides, and consider child genome lists

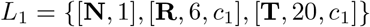

and

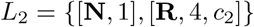

(where *c*_1_ and *c*_2_ are arbitrary branch length elements for these genome list entries) then we need to consider four intersection fragments:

- first, from positions 1 to 3 where both child nodes are of type **N**,
- second, from position 4 to 5 where *b*_1_ is of type **N** and *b*_2_ is **R**,
- third, from position 6 to 19 where both lists are of type **R**,
- fourth, at position 20 where *b*_1_ is **T** and *b*_2_ is **R**.

Calculations for each intersection fragment are performed separately, similarly to how calculations for each site in the Felsenstein pruning algorithm are performed independently. (Note that for datasets with low divergence, the number of intersection fragments will typically be much smaller than the total number of sites.) We describe this process here considering a general non-empty intersection between an entry *e*_1_ of *L*_1_ and an entry *e*_2_ of *L*_2_ — the whole genome list *L*_*n*_ is generated by repeating this process in order of genome position for each non-empty intersection and concatenating the results in *L*_*n*_. For simplicity, we assume that *e*_1_ = [*τ*_1_, *i*_1_, *c*_1_, *v*_1_] and *e*_2_ = [*τ*_2_, *i*_2_, *c*_2_, *v*_2_], that *i* = max(*i*_1_, *i*_2_), and that the intersection fragment between *e*_1_ and *e*_2_ consists of *λ* nucleotides. (If *τ*_1_ = **O** and in other similar cases then we have necessarily *λ* = 1.) Our aim is to calculate the corresponding entry *e* = [*τ, i, l, v*], which refers to the partial likelihoods for the intersection fragment of *λ* nucleotides starting at position i for the internal node *n*; this entry will then be added to genome list *L*_*n*_. Graphical examples of the cases below are given in Extended Data Fig. S6.

- When at least one of *τ*_1_ and *τ*_2_ is **N** (Extended Data Fig. S6B, C), since one child node contributes no information, we need only use the genome list entry information of the other child. If for example *τ*_1_ = **N** we have *e* = [*τ*_2_, *i, c*_2_ + *l*_2_, *v*_2_]. Note however that if also *τ*_2_ = **N** then we don’t need to keep track of the branch length element of *e* (Extended Data Fig. S6B), and if *τ*_2_ ≠ **O** the partial likelihood vector element of e is also unnecessary (Extended Data Fig. S6C).
- If *e*_1_ and *e*_2_ are of the same type *τ*_1_ = *τ*_2_ ∈ {**R**,**A**,**C**,**G**,**T}** (Extended Data Fig. S6D) then any mutational history involving a different nucleotide at the parent node would have considerably lower likelihood, so we define *e* as of type *τ* = *τ*_1_ = *τ*_2_. The branch length entry of *e* is *l* = 0 since type *τ* is considered observed at node *n*, and no partial likelihood vector *v* is required, resulting in *e* = [*τ, i*, 0,].
- If *τ*_1_ ≠ *τ*_2_ and both *τ*_1_, *τ*_2_ ∈ {**R**,**A**,**C**,**G**,**T}** (Extended Data Fig. S6E), the two likelihoods corresponding to these two nucleotides at node *n* will have similar orders of magnitude, and so we set *τ* = **O**. We can assume for simplicity that *τ*_1_ and *τ*_2_ represent individual nucleotides (if, for example, *τ*_1_ = **R** then we can equivalently consider *τ*_1_ as the reference nucleotide at the considered position). We approximate the relative partial likelihoods at *n* (the entries of *v*) as 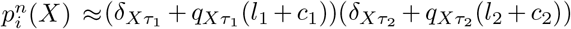 ; here 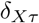 is the Kronecker delta. We then normalize *v*, to obtain entry *e* = [**O**, *i*, 0, *v*].
- The last case is when *τ*_1_ = **O** or *τ*_2_ = **O**. In the most complex case *τ*_1_ = *τ*_2_ = **O** we approximate the partial likelihoods as 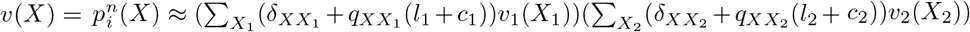 where *v*(*X*) is the entry of *v* corresponding to nucleotide *X*. We then normalize *v*; if only one nucleotide has a value above *ϵ* in *v*, then we set *τ* to this nucleotide (if this nucleotide is the reference nucleotide at site i, we set *τ* = **R**). Therefore *e* = [*τ, i*, 0, *v*], where *v* might be absent in case *τ* ≠ **O**. The case in which only one of *τ*_1_ and *τ*_2_ is **O** is dealt with similarly.

These calculations are iterated over all intersection fragments, which together represent a partition of all genome positions. Entries of genome list *L*_*n*_ are included in order based on position element *i*. If two consecutive entries of *L*_*n*_ are of type **R** and have the same branch length, we merge them into a single entry of type **R**.

The computational demand of this approach is linear in the total number of entries of all the genome lists in the tree, since the maximum computational demand for creating a genome list entry is a constant, no matter the number of sites represented by the entry.

### Other partial likelihoods

So far we have discussed partial likelihoods as in Equation 1. Normally these likelihoods are sufficient for phylogenetic inference. However, when using a non-stationary model, additional types of likelihoods are useful^4^. Here we also use these additional likelihoods and represent them with additional genome lists. Furthermore, for most nodes of the tree, we also calculate genome lists representing relative likelihoods considering all the data in the alignment, which correspond to ancestral state reconstructions^5^. We present the details of these genome lists in Supplementary Methods Section S1.2.

### Phylogenetic placement

Phylogenetic placement is the task of adding a new sequence onto an existing phylogenetic tree^6^. We perform phylogenetic placement within the context of stepwise addition^7^ to construct an initial phylogenetic tree. We start from a tree containing only one sample and iteratively expand it by placing new samples on it one at the time.

First, given a tree *ϕ* and a new sample *s* we look for the region of *ϕ* where to best place *s*. We traverse *ϕ* starting from the root, and we typically do not traverse the whole tree, but instead traverse only a small portion of the internal, terminal, and mid-branch nodes of *ϕ*, stopping traversing into subtrees if the placement at their root looks unpromising (Fig. 2). For each node *n* that we traverse, we use its ancestral state reconstructions and combine them with the genome list representing the partial likelihoods of *s* to obtain the placement score of *s* at *n* (described in detail in Supplementary Methods Section S1.3). As we traverse the tree, we keep track of *B*_*s*_, the best placement score found so far for *s*. If, while traversing the tree, the placement score at an internal node *n* worsens by a certain margin (by default 200 log-likelihood units worse than *B*_*s*_) or at least a certain number of times (by default five times) moving from the direct ancestors of *n* to *n*, then we do not traverse the tree further downward in the subtree of the descendants of *n*. We do not attempt placement at nodes with a branch length of 0 above them, which are part of polytomies.

Once we have identified the node with the best placement likelihood score, we search in detail the exact point of the phylogeny near this node where the new sample is best placed on the tree (Supplementary Methods Section S1.4).

Every time we add a new sample to the tree, we consequently update the genome lists in the tree. Because we consider relative likelihoods, we typically only need to update the genome lists for a small portion of the tree after each new sample placement (Supplementary Methods Section S1.5).

If the genome of a new sample to be placed on the tree is found to be identical or less informative than the genome of a sample already in the tree, we record it as such and only add it to the estimated phylogeny at a later stage (Supplementary Methods Section S1.6).

During estimation of the initial phylogeny by stepwise addition, we also estimate the substitution model (Supplementary Methods Section S1.7), which we then consider fixed in the next stage of MAPLE.

### Tree topology improvement

After estimating an initial tree via stepwise addition, we attempt at improving the topology of the tree using custom subtree pruning and regrafting^7^ (“SPR”) proposals. These work in a very similar way as sample placement, and are described in detail in Supplementary Methods Section S1.8.

### Software implementation

We implemented our methods in a Python3 script available from https://github.com/NicolaDM/MAPLE. For efficiency, we recommend its execution with the pypy3 implementation of Python https://www.pypy.org/#!.

### Other Phylogenetic methods considered

We compare the performance of MAPLE to efficient and popular maximum likelihood phylogenetic methods that are often used to analyse large sequence datasets: IQ-TREE v2.1.3^8^, FastTree v2.1.11^9^ (double precision, no SSE3), and RAxML-NG v1.0.2^10^. For all these methods we adopt a GTR substitution model^11^. We also consider the parsimony-based method matOptimize^12^,a recent approach to improving the accuracy of UShER^13^ trees), which has been tailored for SARS-CoV-2 datasets. We selected program options to permit fair comparison of methods, with each being tuned to the largest problems it could analyze on available hardware. In detail:

We ran IQ-TREE 2 with options “-quiet” to reduce screen output, “-nt 1” to use only one core per replicate on our cluster and “-fast”, with which only nearest neighbour interchange (NNI) moves are used. For simulations with rate variation we used a GTR+G model.

FastTree 2 was executed with options “-quiet” to limit screen output, “-nosupport” to skip support value computations, and “-nocat” to ignore rate variation (except for simulations with rate variation, for which we use “-cat 4”). We also used option “-fastest” to reduce the time demand of NNI steps.

RAxML-NG was run with options “- -threads 1” to use only one core per replicate on our cluster, “- -blmin 0.000005” to increase the minimum branch length considered and “- -tree pars{1}” to start the tree search from a parsimony tree. For simulations with rate variation we used a GTR+G model.

UShER and matOptimize were run with option “-T 1” to utilize a single thread per replicate, and were run using the vcf input file format (option “-v”). matOptimize was run starting from the initial tree estimate of UShER and using option “-n” to avoid the creation of intermediate files.

We ran MAPLE with default parameters and using PyPy (v7.3.5 with GCC 7.3.1 20180303 for Python 3.7.10; see https://www.pypy.org/#!).

Additional options considered for these and additional methods are described in Supplementary Methods Section S1.9, with corresponding results reported in Extended Data Figs. S2–S5.

### Real SARS-CoV-2 sequence data

We randomly subsampled without replacement a given number of sequences from the 540,520 whole genomes that were represented both in the 31 March 2021 global unmasked SARS-CoV-2 alignment from GISAID^14^ and in the corresponding phylogenetic tree (https://www.gisaid.org/). We did not mask sites or filter out sequences. We use the consensus of all the sequences in the global GISAID alignment as reference genome for MAPLE. When measuring running times, we did not consider the cost of creating the input alignment for a given method.

### Simulated SARS-CoV-2 sequence data

For real datasets, we have the drawback of not knowing the true underlying phylogenetic tree, which makes it harder to assess the accuarcy of different phylogenetic inference methods. For this reason, we also simulated SARS-CoV-2 alignments of known phylogeny and substitution dynamics. We used as background “true” tree the publicly available 26 October 2021 global SARS-CoV-2 phylogenetic tree from http://hgdownload.soe.ucsc.edu/goldenPath/wuhCor1/UShER_SARS-CoV-2/^15^ representing the evolutionary relationship of 2,250,054 SARS-CoV-2 genomes as obtained using UShER^13^. We used phastSim v0.0.3^16^ to simulate sequence evolution along this tree according to SARS-CoV-2 non-stationary neutral mutation rates^17^ and using the SARS-CoV-2 Wuhan-Hu-1 genome^18^ as root sequence. We simulated three different scenarios:

- the “basic” simulation scenario (no rate variation and full genomes available)
- the “rate variation” scenario, where we allow different genome positions to evolve at different speeds in our simulations to mimic the effect on genome evolution of variable mutation rates and selective pressures along the genome. We simulated four genome site categories, all with the same frequency and with relative substitution rates of 0.1, 0.5, 1 and 2
- the “sequence ambiguity” scenario, where we modified the simulated sequence data of the basic simulation scenario to include ambiguity characters. To realistically mimic amplicon drop-out effects^19^, for each simulated sequence we sample one random sequence from the real dataset and copy-paste from it the stretches of “N” and gap “-” characters into the simulated sequence. Additionally, because contamination and mixed infections can result in individual ambiguity characters specifically at phylogenetically informative sites of the genome^20^, we count the number of isolated ambiguous characters in the real sequence, and we mask an equal number of randomly selected SNPs (differences with respect to the reference genome) in the simulated sequence. If more isolated ambiguous characters are observed in the real sequence than SNPs in the simulated sequence, then we simply mask all SNPs in the simulated sequence.

### Comparison of methods’ performance

We measured the computational demand of different approaches in estimating phylogenies by tracking the running time and maximum memory demand of all methods. All methods were run in parallel, assigning one thread per replicate per method. Since matOptimize requires an initial run of UShER, the running time of matOptimize is defined as the sum of the time it took to execute UShER followed by matOptimize; the maximum memory demand for matOptimize was defined as the highest of the maximum memory demands of the two methods.

We used two methods to compare the topological inference accuracy of different approaches. The first compares the likelihoods of the estimated tree topologies. Trees with higher topology likelihoods are interpreted as better estimates. Since the phylogenetic likelihood of the same tree measured by different software can differ due to different approximations employed, we use the same software, IQ-TREE 2, to calculate the likelihood of the topologies inferred by all methods. To make the comparison of topological accuracy of different methods even more fair, in particular considering that maximum parsimony methods UShER and matOptimize do not represent branch lengths in the same way as maximum likelihood methods and do not estimate substitution models, when measuring topology tree likelihoods we run IQ-TREE 2 using the tree to be assessed as starting tree, and performing model and branch length optimization but without attempting topological improvements. In simulations with rate variation we run IQ-TREE 2 with a GTR+G model with four categories; otherwise we use a plain GTR model. Note that the use of IQ-TREE 2 for tree topology likelihood estimation limits the size of the trees that can be assessed due to the memory demand of the software.

The second measurement of phylogenetic accuracy (only available for simulated data for which the correct tree is known) is to calculate the Robinson-Foulds distance^21^ between an inferred tree and the corresponding true simulated tree. This distance gives a measure of how topologically close an inferred tree is to the true tree, and therefore quantifies inference error. We consider trees as unrooted, collapse all branches on which no simulated mutation events occurred, and collapse all branches shorter than a minimum branch length (defined by the minimum branch length considered by each estimation method) so as to represent trees as multifurcating when a method finds little or no support for the local branching order. Robinson-Foulds distance calculations were performed with a custom implementation of Day’s algorithm^22^.

## Data availability

All real data used in this manuscript is available from the GISAID^14^ (https://www.gisaid.org/) 31 March 2021 which requires acceptance of the GISAID data sharing conditions.

## Code availability

The code is available from https://github.com/NicolaDM/MAPLE.

## End notes

## Acknowledgements

N.G. and N.D.M. were supported by the European Molecular Biology Laboratory (EMBL). Y.T. was supported by the Centers for Disease Control and Prevention grant BAA 200-2021-11554. R.C.-D. was supported by funding from the Schmidt Futures Foundation, by an Alfred P. Sloan foundation fellowship, and by NIH/NIGMS grant R35GM128932. B.Q.M. was supported by a Chan-Zuckberg Initiative grant for essential open source software. The funders had no role in study design, data collection and analysis, decision to publish, or preparation of the manuscript. We are very grateful to GISAID and all the groups who shared their sequencing data. A full list of these is available from https://github.com/roblanf/sarscov2phylo/tree/master/acknowledgements.

## Author contributions

N.D.M. conceived and implemented the methods, performed the simulations and real data analyses, and wrote the first draft of the manuscript.

N.G. supervised the work and finalized the manuscript.

B.Q.M., R.C.-D., Y.T. and P.K. provided support during the analyses, method implementation and the drafting of the manuscript.

