## Supplement for "Maximum likelihood pandemic-scale phylogenetics"

### Extended Data Figures

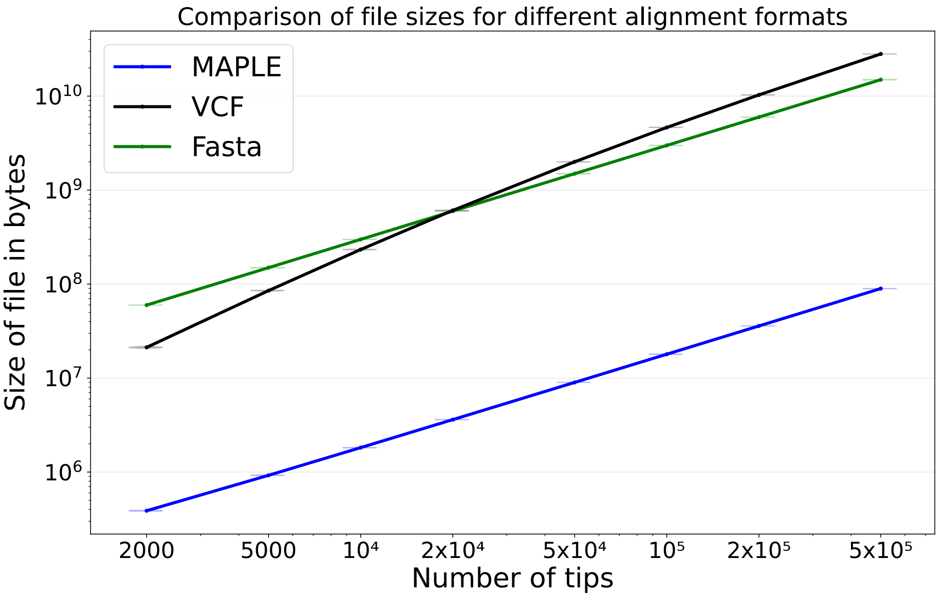


Extended Data Figure S1: **Comparison of file sizes of SARS-CoV-2 genome alignments using different alignment formats**. On the Y axis on a logarithmic scale we show the sizes of alignment files for each format considered, expressed in bytes. On the X axis is the number of sequences in the dataset considered on a logarithmic scale. Here we consider random subsamples of our real SARS-COV-2 alignment data. Violin plots (often variation within one plot is not visible, collapsing the violin plots into horizontal lines) summarize values for 20 replicates, and dots represent their mean.

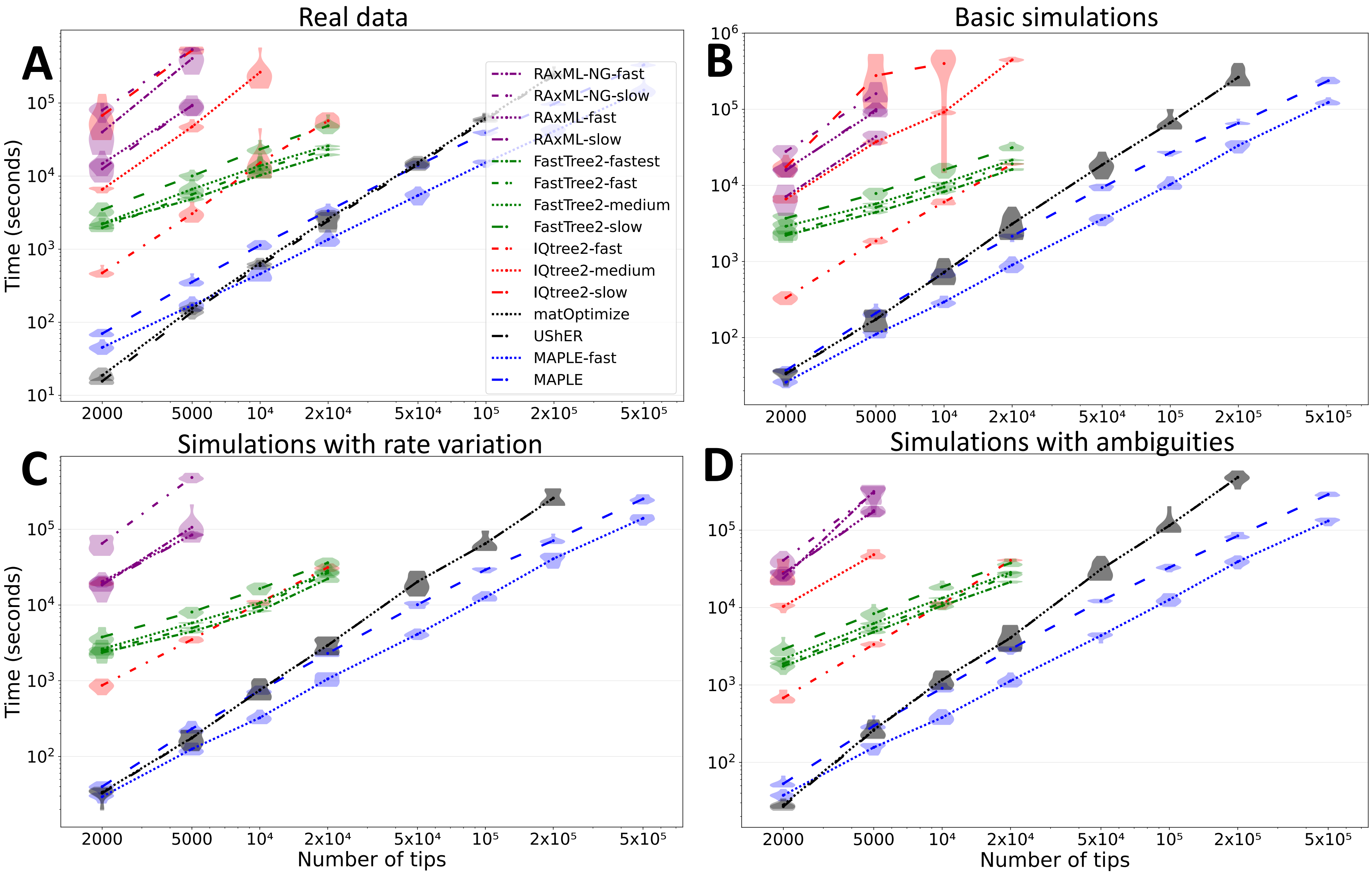


Extended Data Figure S2: **Comparison of running times of all considered methods and options for phylogenetic inference from SARS-CoV-2 genomes**. On the Y axis on a logarithmic scale we show the number of seconds it takes to run each method. On the X axis is the number of sequences in the dataset considered on a logarithmic scale. Different line styles and colors represent different options for each method, as denoted in the legend and Section S1.9. We ran each method and set of options up to the maximum dataset size that was achievable due to time and memory limitations. Violin plots summarize values for 10 replicates, and dots represent their mean. **A** Results for subsamples from the real SARS-CoV-2 dataset. **B** Simulated datasets with no rate variation or ambiguity. **C** Results on simulated data with rate variation but no ambiguities. **D** Simulated data with sequence ambiguities but no rate variation.

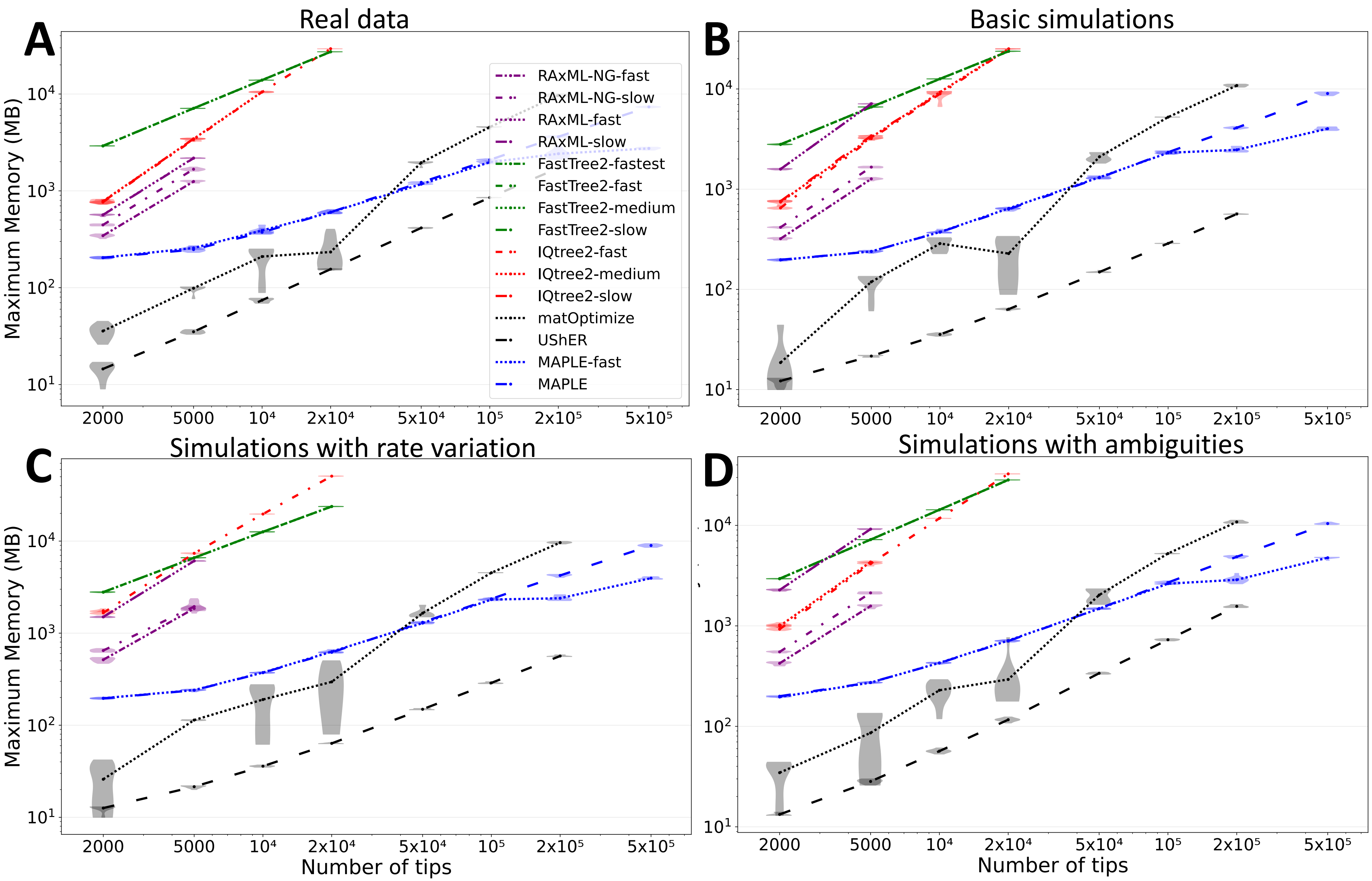


Extended Data Figure S3: **Comparison of maximum memory demand of all considered methods and options for phylogenetic inference from SARS-CoV-2 genomes**. On the Y axis on a logarithmic scale we show the maximum RAM memory demand in MB required to run each method. Other details are the same as in Extended Data Fig. S2.

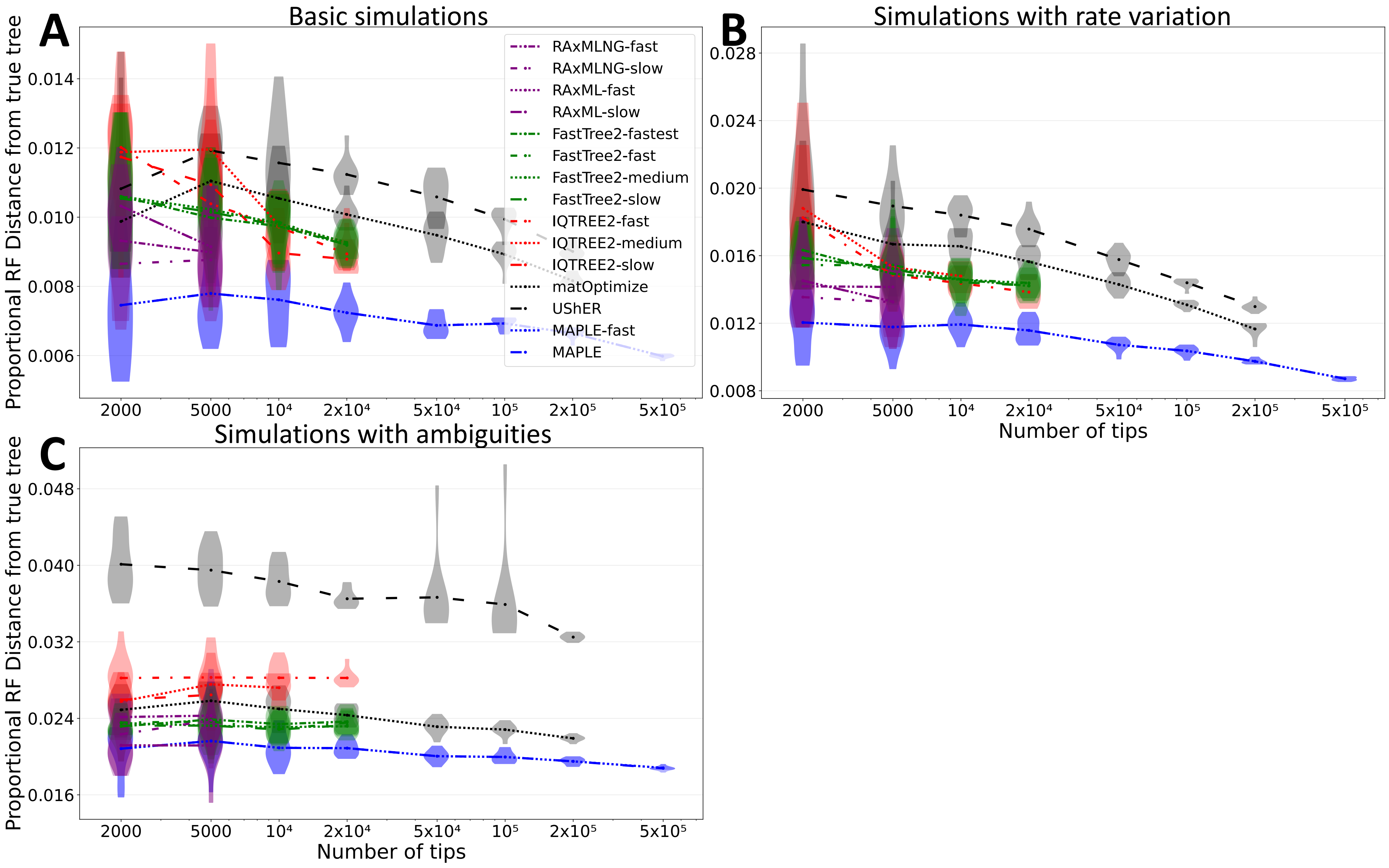


Extended Data Figure S4: **Comparison of proportional Robinson-Foulds distances of inferred trees from the correct simulated trees**. On the Y axis we show the proportional Robinson-Foulds distances (that is, normalized by $2(m-3)$ with $m$ the number of samples in the tree) of the tree estimated by each method with respect with the true simulated tree of the corresponding scenario and replicate. We collapsed tree branches of the simulated trees where no mutation event was simulated. Trees were compared as unrooted, and polytomies were compared as such (we collapsed branches of inferred trees with length equal to the minimum allowed length by the corresponding inference method). **A** Results for simulated datasets with no rate variation or ambiguity. **B** Results on simulated data with rate variation but no ambiguities. **C** Simulated data with no rate variation but with ambiguities. Other details are the same as in Extended Data Fig. S2.

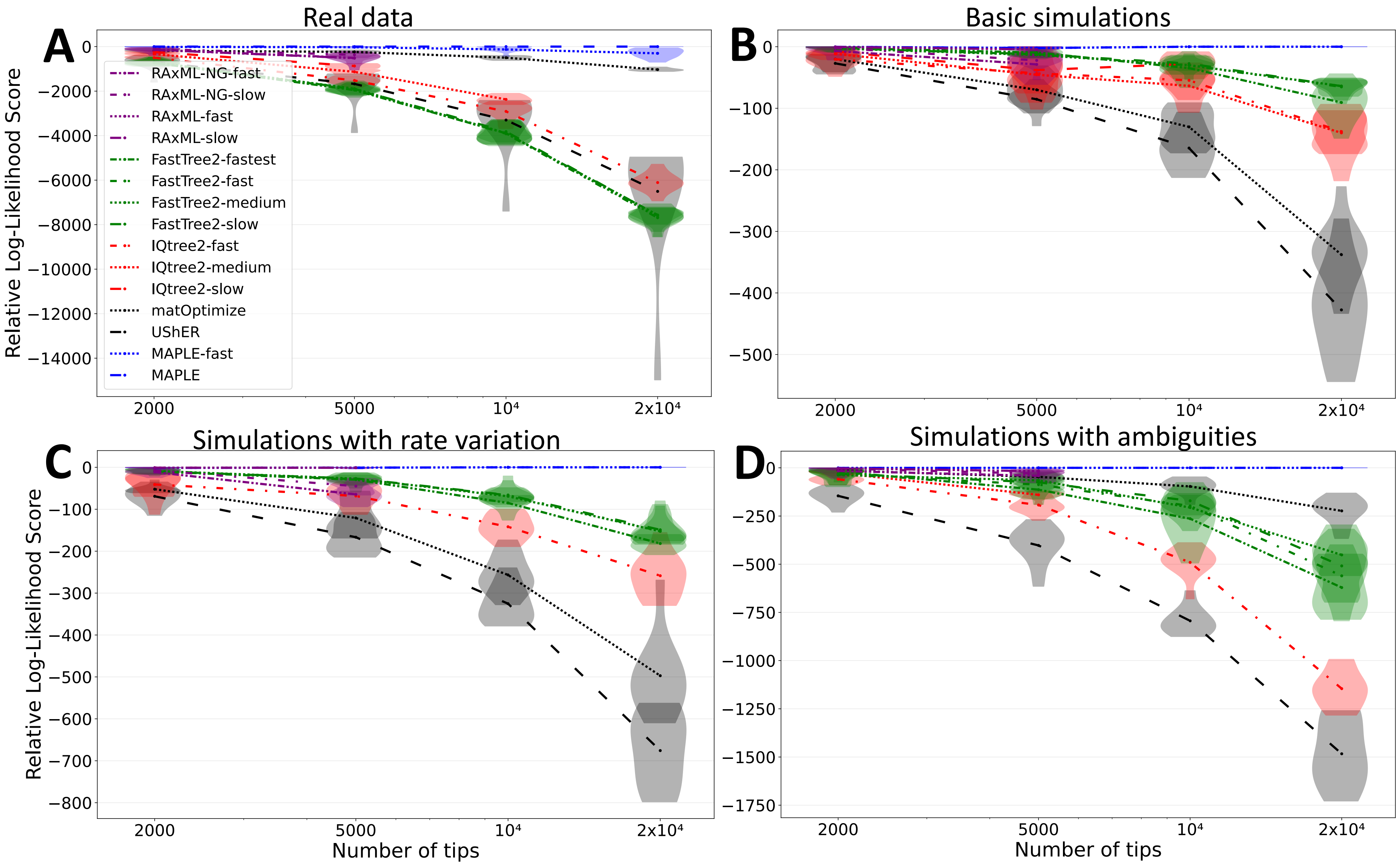


Extended Data Figure S5: **Comparison of relative likelihood scores of trees inferred by different phylogenetic methods from SARS-CoV-2 genomes**. MAPLE leads to more accurate tree reconstruction (tree topologies with higher likelihoods) both in real data and simulations. On the Y axis we show the relative log-likelihood scores of the tree estimated by each method, as in Fig. 3F, with higher scores representing more likely tree estimates. Other details are the same as in Extended Data Fig. S2.

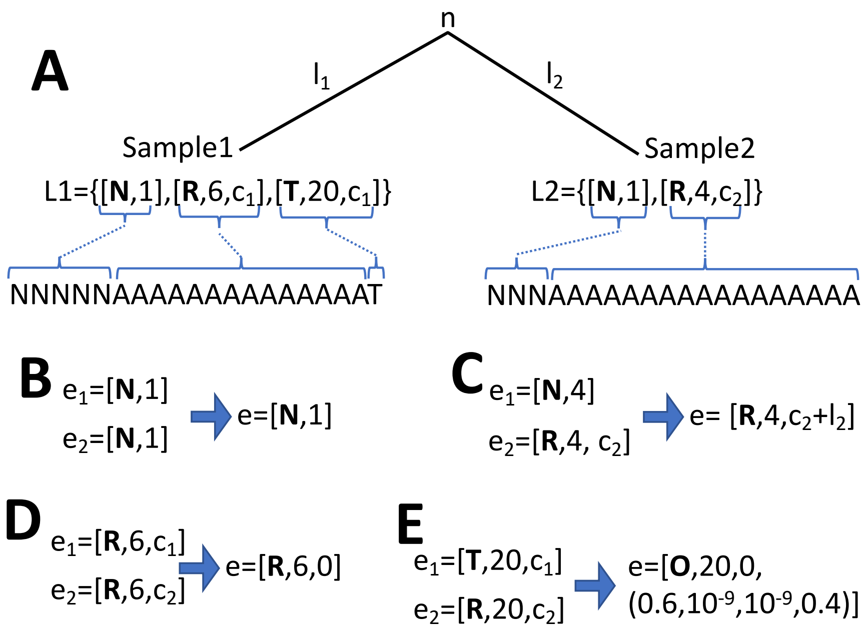


Extended Data Figure S6: **Graphical example of the merging of genome lists**. Here we consider the example genomes considered in the text, with a reference genome consisting of 20 "A" nucleotides, partial likelihood genome lists $L_1$ and $L_2$, and a phylogeny with two branches, one of length $l_1$ leading to Sample1 and genome list $L_1$, and one of length $l_2$ leading to Sample2 and genome list $L_2$. **A** Graphical representation of the phylogeny, of the two observed genomes and of the partial likelihood genome lists at the tips. Blue parentheses and lines highlight the correspondence between genome list entries and portions of the observed genomes at the tips of the tree. Our aim here is to calculate the partial likelihood genome list for the most recent common ancestor $n$ of Sample1 and Sample2. Parameters $c_1$ and $c_2$ here usually would have value 0 since they refer to tips of the tree - however, for internal nodes these values could be strictly positive, so we use general parameter names here to give more generality. **B** For the first intersection fragment of the genome consisting of the first three positions, both children node partials contain no information (they are of type **N**), so the same is true for their parent (which will also be of type **N**). **C** For positions 4 and 5, Sample1 provides no information while Sample2 presents the reference allele. In this case the parent node genome list entry will be of type **R**, but we also keep track of the branch length $l_2$ using the branch length element of the entry. This way, the branch length element at $n$ now records the evolutionary distance $(c_2 + l_2)$ between $n$ and the last visited node in the tree with no state uncertainty (Sample2). **D** From positions 6 to 19 both child node genome list entries are of type **R**, and so the same is true for the parent node at their intersection fragment. The corresponding parent node genome list entry will have branch length element set to 0, which is the same as considering the reference alleles observed exactly at the parent node. **E** At the last position of the genome, while at Sample1 we observe nucleotide "T", at Sample2 we observe nucleotide reference "A"; in this case, the parent node genome list entry will be of type **O**, and we calculate an explicit partial likelihood vector with the relative likelihoods of all four nucleotides. The branch length element of the genome list entry is set to 0, since the relative partial likelihoods refer to the parent node.

### S1 Supplementary Methods

Our approach to tree inference differs from traditional maximum likelihood ("ML") phylogenetic approaches in that we concisely represent genetic sequences and partial likelihood vectors (see Main Text Methods section); we not only make use of these concise representations to reduce memory demand, but we also use novel algorithms to efficiently calculate and update likelihoods. To further reduce time complexity in likelihood calculations, and to allow non-stationary substitution models (which better describe SARS-CoV-2 evolution[1]) we adapt the strategy of Boussau and Gouy[2] to our scenario, storing partial likelihoods from different subsets of the data at each node, as described in Section S1.2.

Our first step in phylogenetic inference is to use ML stepwise addition[3] to build an initial tree from scratch, given only the input genetic data. This means that, starting from a tree containing just one sample, we iterative add (or "place") new samples to the current tree one at the time, so that at each step the tree grows in size by one sample. We aim to do this both efficiently and accurately, so that the initial tree itself already represents a reasonable phylogenetic estimate. In fact, at each step, we place a new sample on the tree so to minimize the likelihood cost of the addition, but at the same time avoiding the traversal of the full phylogeny. At the same time as we build the initial tree, we also estimate the substitution process (the model of sequence evolution). The process of building the initial tree is described in more detail in the Main Text Methods section, and the way that the placement score is calculated is described in Section S1.3.

Once we finalize the initial phylogenetic tree by stepwise addition, we then attempt at improving the tree by proposing changes to its topology and branch lengths. We use SPR (Subtree Prune and Regraft) moves to change the tree topology, similar to other phylogenetic methods, but instead of limiting the search radius of the SPR moves in terms of number of branches separating the initial and proposed placement of subtree, as typically done, we instead use an approach based on likelihood thresholds that combines efficiency with accuracy; for more details, see Section S1.8. After a number of series of these tree updates (with number specified by the user), the tree obtained represents the ML tree estimate of MAPLE.

Most of the symbols and expressions used throughout here and the Main Text Methods section are summarized in Table S1.

#### S1.1 Formal definition of genome list entry intersections

Give two genome lists $L_1$ and $L_2$, assume that entry $e_1$ of $L_1$ has position element $i_1$ and "ends" at position $q_1$ (meaning that $q_1 + 1$ is the position element of the next entry in $L_1$, or that $e_1$ is the last element of $L_1$ and $q_1$ is the length of the reference); similarly, assume that entry $e_2$ of $L_2$ has position element $i_2$ and "ends" at position $q_2$. The intersection of these two entries will be non-empty if and only if $q_1 \geq i_2$ and $q_2 \geq i_1$. If this is the case, the intersection genome list $L$ of $L_1$ and $L_2$ will contain an entry $e$ corresponding to the intersection segment of $e_1$ and $e_2$, which will have starting position $\max(i_1, i_2)$ and end position $\min(q_1, q_2)$. $L$ will contain all such entries resulting from non-empty intersections of entries of $L_1$ with entries of $L_2$. See Extended Data Fig. S6A for a graphical representation.

| Expression | Meaning |
|---|---|
| $\phi$ | Phylogenetic tree |
| $n$ | Generic node of $\phi$ |
| $L$ | Genome length (number of alignment columns) |
| $i$ | Generic genome position $1 \leq i \leq L$ |
| $A$ | Genetic data (multiple sequence alignment) |
| $A_i$ | Column $i$ of alignment $A$ |
| $A_i^n$ | Sub-vector of $A_i$ corresponding to all the descendants of $n$ |
| $M$ | Model of sequence evolution |
| $X$ | Generic nucleotide |
| $p_i^n(X)$ | Partial likelihood at node $n$ and position $i$ conditional on nucleotide $X$: $p(A_i^n \mid X, M, \phi)$ |
| $\tilde{p}_i^n(X)$ | Relative (normalized) likelihood: $p_i^n(X)/\sum_D p_i^n(D)$ |
| $e$ | Generic genome list entry $e = (T, i, l, v)$ |
| $\tau$ | Generic genome list entry type $\in \{\mathbf{A},\mathbf{C},\mathbf{G},\mathbf{T},\mathbf{R},\mathbf{N},\mathbf{O}\}$ |
| $l$ | Evolutionary distance from the node of calculation of the partial likelihoods represented by the genome list entry |
| $v$ | Generic vector of partials $(\tilde{p}_i^n(X))_X$ |
| $\epsilon$ | Lower threshold of negligibility for relative likelihoods $\tilde{p}_i^n(X)$ |
| $K$ | Total likelihood, tracks the likelihood contributions across the genome and the subtree of the considered node |
| $L_n$ | Genome list at node $n$ |
| $t(i)$ | Cumulative substitution rate up to reference position $i$ |
| $\pi(X)$ | Root frequency of nucleotide $x$ |
| $Q$ | The substitution rate matrix |
| $q_{X_1X_2}$ | Substitution rate from nucleotide $X_1$ to $X_2$ |
| $-q_{XX}$ | Total substitution rate from nucleotide $X$ |
| $r_i$ | Nucleotide at position $i$ of the reference genome |
| $A_i^{n\uparrow}$ | Data at position $i$ for the tree except the descendants of $n$ |
| $p_i^{n\uparrow\leftarrow}(X)$ | Likelihood of the data from left child and parent node |
| $p_i^{n\uparrow\rightarrow}(X)$ | Likelihood of the data from right child and parent node |
| $p_i^{n\uparrow\rightarrow\leftarrow}(X)$ | Likelihood for all data |
| $s$ | Generic sample |

Table S1: Explanation of main symbols and expressions used in the Methods section.

#### S1.2 Other partial likelihoods

So far we have discussed partial likelihoods of the form discussed in Equation 1. Normally these likelihoods are sufficient for phylogenetic inference. However, when using a non-stationary model, additional types of likelihoods are useful[2]. We follow this same approach here, adapted however to our concise likelihood representation.

Each internal node $n$ of our binary tree $\phi$, with the exception of the root, is connected to three other nodes: two children $b_1$ (the left child) and $b_2$ (the right child) and the parent node $P$. The partial likelihoods of the previous section, $p_i^n(X)$ (which we will refer to here as "lower likelihood"), can be considered as the likelihood of the data "arriving to $n$" from $b_1$ and $b_2$, that is the likelihood considering the data of all descendants of $b_1$ and $b_2$.

In many circumstances, however, for example when we want to evaluate the likelihood score of adding a new sample to the tree as a descendant of a node $n$ (discussed in section S1.3), or the likelihood score of removing a subtree and re-grafting it as a descendant of $n$ (section S1.8), we need to consider all of the information in the tree and alignment. To do so efficiently it is convenient to have available, at a node $n$, pre-computed likelihoods that account for all of the data. These "overall likelihoods" are:

$$p_i^{n\uparrow\rightarrow\leftarrow}(X) = p(A_i, X|M, \phi) \tag{3}$$

where $A_i$ like before is all the data in the alignment at site $i$, and $M$ is the sequence evolution model; here we use arrow $\uparrow$ to represent the fact that we consider the data "arriving" at $n$ from its parent node, and similarly arrows $\rightarrow$ and $\leftarrow$ referring to the right and left child nodes of $n$, when these exist. These overall likelihoods $p_i^{n\uparrow\rightarrow\leftarrow}(X)$ can be approximately calculated and concisely represented similarly to lower likelihoods; again, we only keep track of relative likelihoods, so in practice we only record normalized values $\tilde{p}_i^{n\uparrow\rightarrow\leftarrow}(X)$ corresponding to the posterior probabilities nucleotides at node $n$ and site $i$, that is, they represent the ancestral state reconstructions[4].

In addition to calculating overall likelihoods genome lists for each node of the tree (either internal or terminal), we also calculate them for branch mid-points at all non-zero length branches; these lists will help us efficiently and accurately perform tree exploration in the following sections.

To calculate overall likelihoods for the root node we need to consider the root frequencies of the nucleotides. The overall likelihoods at the root are the lower likelihoods multiplied by the root nucleotide frequencies: $p_i^{n\uparrow\rightarrow\leftarrow}(X) = \pi(X)p_i^n(X)$, where $\pi(X)$ is the root frequency of nucleotide $X$. Overall likelihood genome lists can similarly be obtained from lower likelihood genome lists.

Overall likelihood genome lists at non-root nodes of the tree are instead less straightforward to calculate, and to do it efficiently, we define and keep track of two additional sets of likelihoods (corresponding to two additional genome lists) at each node of the tree. The "upper-left" likelihood $p_i^{n\uparrow\leftarrow}(X)$ is defined as the likelihood of the data that is "passed on" to $n$ from its parent node $P$ and its left child $b_1$. To formally define this likelihood, we call $A_i^{n\uparrow}$ all the data in the alignment column $A_i$ that does not represent observations for any descendant of $n$, so containing all data in $A_i$ that is not found in $A_i^n$. The upper-left likelihood is defined as

$$p_i^{n\uparrow\leftarrow}(X) = p(A_i^{b_1}, A_i^{n\uparrow}, X|M, \phi), \tag{4}$$

while similarly the upper-right likelihood is defined as

$$p_i^{n\uparrow\rightarrow}(X) = p(A_i^{b_2}, A_i^{n\uparrow}, X|M, \phi). \quad (5)$$

For the root node, given the fact that it does not possess a parent node, its upper-left (respectively, upper-right) likelihood is calculated combining the lower likelihoods of its left child $p(A_i^{b_1}|X, M, \phi)$ (respectively, right child $p(A_i^{b_2}|X, M, \phi)$) with the root nucleotide frequencies, as done for the overall likelihoods of the root. For all other nodes, instead, we need to combine likelihood vectors using a very similar approach to the algorithm described in the Main Text Methods. If $n$ is a left (right) child of $P$, to calculate $p_i^{n\uparrow\leftarrow}(X)$, we need to combine the upper-right (upper-left) likelihoods of $P$ with the lower likelihoods of $b_1$, and similarly for $p_i^{n\uparrow\rightarrow}(X)$. Finally, to calculate the overall likelihoods at $n$ we can use different combinations, for example we can combine the upper-right likelihoods at $n$ with the lower likelihoods at $b_1$.

In addition to calculating overall likelihood genome lists at internal nodes of the tree, we also calculate them at terminal nodes of the tree (corresponding to samples) and at some mid-branch nodes (nodes that we add in the middle of branches that have length beyond a certain threshold). We create these additional overall likelihood genome lists so to also allow efficient placement of new sample near samples already in the tree and as descendants of mid-branch nodes. if a terminal node is the left child of its parent, then its overall likelihood genome list is calculated by combining its lower likelihood with the upper-right likelihood genome list of its parent; similarly if the node is a right child. For mid-branch nodes, again, if the node at the lower end of the branch is the left child of its parent, then we combine the upper-right likelihood genome list of the parent with the lower likelihood of the child; similarly if the node at the lower end of the branch is a right child.

#### S1.3 Phylogenetic placement scoring

When looking for the best placement of a new sample $s$ to an existing tree $\phi$, for each node $n$ we traverse, we use its overall likelihood genome list (representing the relative likelihoods $\tilde{p}_i^{n\uparrow\rightarrow\leftarrow}(X)$) and combine it with the lower likelihood genome list of $s$ (which represents the $\tilde{p}_i^s(X)$ relative likelihoods) to obtain the placement score of $s$ at $n$; this is done in a way similar to our algorithm for calculating genome lists described in the Main Text Methods, with the simplifications that:

- Node $n$ is assumed to be ancestral to $s$.
- We don't need to calculate a genome list resulting from the merging, but only a relative likelihood cost for adding the new sample to the tree at the given location in the tree.

Just like in the Main Text Methods, we describe this process here by calculation the contribution to the log-likelihood score of a general non-empty intersection between an entry $e_1$ of $L_1$ (the overall likelihood genome list of the node $n$ of the current tree where the new branch and sample are placed), and an entry $e_2$ of $L_2$ (the partial likelihood genome list of the new sample $s$). The whole log-likelihood score is obtained by summing the log-likelihood contributions of each such non-empty intersection.

For simplicity, we assume that $e_1 = [\tau_1, i_1, c_1, v_1]$ and $e_2 = [\tau_2, i_2, 0, v_2]$, that $i = \max(i_1, i_2)$, and that the intersection fragment between $e_1$ and $e_2$ consists of $\lambda$ nucleotides, that is $\lambda = \min(q_1, q_2) + 1 - i$ (where as before $q_1$ is the last genome position represented by $e_1$ and $q_2$ is the last genome position represented by $e_2$); in case $\tau_1 = \mathbf{O}$ and other similar cases then we have necessarily $\lambda = 1$. We also assume that the length of the new placement branch added to the tree is $l_2$.

- When at least one of $\tau_1$ and $\tau_2$ is **N**, the fragment does not contribute to the likelihood score.
- When $e_1$ and $e_2$ are of the same type: $\tau_1 = \tau_2 \in \{\mathbf{R},\mathbf{A},\mathbf{C},\mathbf{G},\mathbf{T}\}$, the two children nodes of $n$ support the same nucleotide with negligible uncertainty. Since the evolutionary distances separating the nodes of the tree are assumed to be short, and therefore any mutational history involving a different nucleotide at the parent node would not be parsimonious and would have considerably lower likelihood, then the contribution of this fragment to the likelihood score is the probability of the mutational history with no events at the considered genome positions and tree branch; for example, if $\tau = \mathbf{A}$, the log-probability that no mutation event happened along the evolutionary distance $l_2 + c_1$ is approximated as $\log(1 + (l_2 + c_1)q_{AA}) \approx (l_2 + c_1)q_{AA}$. The same approach is taken for $\tau$ equal to **C**, **G** or **T**. Notice that this step does not require the computationally demanding calculation of logarithms. The scenario $\tau = \mathbf{R}$ works similarly, except that this time we have to consider the log-probability contribution over all $\lambda$ sites of the considered fragment. This is done efficiently by pre-computing the total substitution rate for any prefix stretch of the reference genome $t(i) = \sum_{j=1}^{i} q_{r_j r_j}$ (with $t(0) = 0$ by definition), where $r_j$ is the nucleotide at position $j$ of the reference genome. Then, the total substitution rate for a stretch of $\lambda$ sites from position $i$ is $t(i + \lambda - 1) - t(i - 1)$ and the approximate log-likelihood contribution for the fragment is calculated in constant time as $(l_2 + c_1)(t(i + \lambda - 1) - t(i - 1))$.
- When $\tau_1 \neq \tau_2$ and both $\tau_1, \tau_2 \in \{\mathbf{R},\mathbf{A},\mathbf{C},\mathbf{G},\mathbf{T}\}$, the two children nodes of $n$ support two different nucleotides, and we necessarily have $\lambda = 1$. We can assume for simplicity that $\tau_1$ and $\tau_2$ represent individual nucleotides (that is, if for example $\tau_1 = \mathbf{R}$, then we can equivalently consider $\tau_1$ as the reference nucleotide at the considered position). The log-likelihood contribution of the fragment is then approximated as $\log(q_{\tau_1 \tau_2}(l_2 + c_1))$.
- The last case is when $\tau_1 = \mathbf{O}$ or $\tau_2 = \mathbf{O}$. In this case, at least one of the child nodes of $n$ has likelihoods not concentrated at a single nucleotide (that is, is of type **O**). Here we show as an example likelihood calculation of the most complex case $\tau_1 = \tau_2 = \mathbf{O}$. We approximate the log-likelihood contribution of the fragment as the logarithm of $\sum_{X_1, X_2} v_1(X_1)(\delta_{X_1 X_2} + q_{X_1 X_2}(l_2 + c_1))v_2(X_2)$.

These calculations are iterated over all intersection fragments, which together represent a partition of all genome positions, and we sum up all log-likelihood contributions of the intersection fragment to obtain the log-likelihood score of the placement at node $n$.

#### S1.4 Zooming in on the phylogenetic neighborhood to finalize sample placement

Once we have identified the node (or mid-branch point) in the phylogeny with the best placement likelihood score, we need to identify exactly the point of the phylogeny near this node (or mid-branch point) where the new branch should be attached to the tree, and we need to define the length of this branch. Here we describe how these choices are made based on ML.

If the best placement score was found at a mid-branch point, we only consider different possible placements along this branch. Assuming that the preliminary placement is on a branch with length $l$, the exact point of the preliminary placement will be at height $l/2$ along this branch. First, we try to change this height to $l/4$, and if this leads to a placement likelihood improvement, we further attempt height $l/8$, and so on, until we reach below a certain minimum height (by default $0.2/L$). if the likelihood at height $l/4$ is worse than at $l/2$, then we also attempt at moving the placement upward to height $3l/4$, and if this results in a placement likelihood improvement, we move further up to $7l/8$, and so on, until a certain maximum height is reached (by default $l - 0.2/L$). Every time we attempt placement at a new height we need to calculate a new overall likelihood genome list for the existing phylogeny at the new height; this can be done efficiently using the existing genome lists and the nodes above and below the considered branch. Then, we optimize the length of the new branch added to the tree, by similarly attempting at halving or doubling its length up to a minimum (by default $0.2/L$, but a length of exactly 0 is also attempted) or maximum (by default $40/L$) value.

If the best preliminary placement score is instead at a node $n$, we try to place the sample above $n$ (if the branch above $n$ has length $l$, we attempt heights $< l/2$ in a procedure similar to above) and below $n$, that is, on any branch leading to any child of $n$. If $n$ represents a polytomy (meaning that at least one of the branches directly below $n$ has length 0, which is how we represent polytomy within a formally binary tree), we consider all children of $n$ to be part of this polytomy. If a child of $n$ has a branch above it of length $l$, we only attempt placements at heights $> l/2$ similarly to before. As before, we also optimize the length of the new branch leading to $s$ added to the tree.

#### S1.5 Updating genome lists

Every time we add a new sample to the tree, we need to update the genome lists (representing relative partial lower likelihoods, total likelihoods, upper left likelihoods, and upper right likelihoods) for the nodes of the tree. Here we describe how this can be done efficiently by only traversing a small portion of the tree each after each new sample placement.

We start a tree traversal from the location of the placement, and update genome lists for the node of the placement and the nodes just above and below it. These updated genome lists are then "passed on" to neighbouring nodes, following the direction in which the tree is traversed. If at any step, the updated likelihoods are identical to the old ones (up to the threshold $\epsilon$), making the update unnecessary, the tree traversal in this direction is halted.

For example, assume that a new sample $s$ is added to the tree by placing it on the branch above node $n$ — this means that now $n$ has a new parent node, $P$, of which $n$ is the left child and $s$ is the right child. We first calculate all the genome lists for $P$ using the existing genome lists in the tree and using the lower likelihood genome list for $s$; then we calculate the overall likelihood genome list for $s$; and then we need to update the genome lists of $n$ and all of its descendants. To do this, we pass to $n$ the upper-right likelihood genome list of $P$ and we combine it with the lower likelihood genome list of $n$ to calculate the new overall likelihood genome list for $n$. We similarly update the upper-right and upper-left likelihood genome lists of $n$. If the overall likelihood genome list of $n$ has not changed in this process, no further updates are performed for the genome lists of the descendants of $n$; otherwise, we pass the new upper-right likelihood genome list of $n$ to its left child, and its upper-left likelihood genome list to its right child, and repeat this process for both children. After the update of the genome lists of the descendants of $n$ is completed, we proceed similarly in updating the likelihoods of the nodes that are not descendants of the new node $P$.

#### S1.6 Dealing with less informative sequences

Here we describe an approach that we use to reduce the complexity of the phylogenetic tree: we remove from the tree samples that are identical or less informative than other samples already in the tree.

When placing a sample $s_1$, if we find that its sequence is identical to another one associated with sample $s_2$ already in the tree, instead of adding $s_1$ as its own independent tip to the existing tree, we add it to a specific list of samples identical to $s_2$. This is because a ML placement of $s_1$ is the one that places $s_1$ and $s_2$ exactly at the same spot of the tree, with the two samples separated only by branches of length 0. This is a common approach in ML phylogenetics, where only one representative for each set of identical sequences is considered during phylogenetic inference. Then, after the phylogenetic tree relating the representatives is estimated, all samples previously excluded are added to the tree at the same spot as their corresponding representatives to produce the final tree[5]. We take the same approach here.

However, we go one step further, and not only retain only one representative for each set of identical sequences, but also remove sequences that are less informative than others already added to the tree. What we mean by $s_2$ being less informative than $s_1$ is that $s_2$ and $s_1$ coincide everywhere in their sequence except for positions where $s_1$ is strictly less ambiguous than $s_2$ and not in contradiction with it; for example, $s_2$ might have a "N" character at some position where $s_1$ has a nucleotide letter or any IUPAC ambiguity code. Another example is when $s_2$ has ambiguity character "Y" (meaning "C" or "T"); in this case $s_1$ is more or equally informative than $s_2$ if it has a "C", "T", or "Y" entry at this position.

A more precise description of this approach and an explanation of why it works is given below in Section S1.6.1.

During the placement of $s_2$, if we visit a sample $s_1$ that has an equally or more informative sequence than $s_2$, we add $s_2$ to the specific list of sequences that $s_1$ represents, halt the placement search for $s_2$, and do not extend the tree to include $s_2$. Then, at the end of the tree inference procedure, we add $s_2$ back to the tree at the same place on the tree as $s_1$.

This approach of dealing with less informative sequences is not only useful here, but is applicable more generally to most phylogenetic inference frameworks, and we propose it as a more efficient extension to the typical approach for dealing with identical sequences.

In order to take full advantage of this procedure, before placing samples on the tree one at the time, we sort them based on their number of ambiguous characters and on the number of differences with respect to the reference. This increases the number of times in which a more informative sequence is placed on the tree before a less informative one, so that we can remove more samples from the phylogenetic inference process and reduce overall computational demand. A more thorough comparison of each pair of samples would remove more sequences, but to be performed would require quadratic time in the number of samples.

A caveat of this procedure is that multiple ML trees might exist for the same dataset. For example, if we consider the following three sequences, each 2 bp long: AA, AC, AN, then the tree ((AC:0,AN:0):$l$,AA:$l$) will have, under a reversible substitution model, the same likelihood as the tree ((AA:0,AN:0):$l$,AC:$l$). The approach of removing samples from a tree, depending on how is specifically performed, can introduce biases with respect the selection of one of such trees versus another. If our aim is to estimate any ML tree, then this is not an issue. However, if one also aims to represent the uncertainty in ML inference, for example as in phylogenetic bootstrap[6], then this approach can result in biases if for a sequence one more informative representative is selected more often than other possible alternative; the same is true for our approach and for the traditional approach of removing strictly identical sequences.

##### S1.6.1 Formal definition of sequence informativeness

We will represent the fact that two aligned sequences $a$ and $b$ are identical as "$a = b$", and the fact that $a$ is strictly (non-strictly) more informative than $b$ as "$a > b$" ("$a \geq b$"). What we mean by $a$ being "more informative" than $b$, is that any allele that is possibly present at a position of the genome of the sample of $a$, could possibly also be present at the same position in $b$. In the simple case of an alignment made of a single column, we have that A>N, while A=A, but A and C are not comparable (A$\ngeq$C and C$\ngeq$A). For the following, gaps are treated as the same as ambiguous "N" characters. The other relations for other IUPAC ambiguity codes follow similarly, for example, since Y represents "C or T", we have that C>Y, that Y>N, and that Y and A are not comparable. For longer alignments/sequences, we can simply say that for sequences $a$ and $b$ we have $a \geq b$ if and only if $\forall i, a_i \geq b_i$ where $a_i$ is the $i$th character of sequence $a$ (or more precisely the character of $a$ in alignment column $i$). Under these definitions, the sequences within an alignment form a partially ordered set.

Our sample removal strategy is motivated by the following fact:

**Lemma S1.1.** *If $a \geq b$, then the tree obtained removing $b$ from the alignment, estimating a ML tree, and re-adding $b$ to form a 0-branch-length clade with $a$, is still a ML tree.*

*Proof.* Let's call $T_{-b}$ the ML tree obtained without $b$. The tree obtained by adding $b$ to it will be $T_{-b+b}$, and we have that $T_{-b}$ and $T_{-b+b}$ have the same likelihood, since one can think of $b$ as a descendants of $a$ with 0 distance from it, and no substitutions from it. Now, let's assume *ad absurdum* that a tree $T$ exists for the whole alignment with a higher likelihood than $T_{-b+b}$. We can then remove $b$ from $T$ and obtain a tree $T - b$ with the same or higher likelihood than $T$. This means that $T - b$ has a higher likelihood than $T_{-b}$, which is absurd since $T_{-b}$ was a ML tree, and which proves the lemma. $\square$

Lemma S1.1 can of course be generalised to any number of sequences: if we have $B = \{b_1 \dots b_n\}$ a set of sequences that we remove from the alignment, and $\forall i, b_i \leq a_i$ for some $a_i$ not removed from the alignment, then we can remove all the sequences in $B$, estimate a tree, and then re-add $B$ to the tree "attaching" each to $b_i$ to the corresponding $a_i$, and we would still have a ML tree - which can be proven by sequentially applying lemma S1.1.

This means that, in order to infer a ML tree, we only need to infer a ML tree for the maximal sequences (the sequences that have no other sequence more informative than them) in the alignment, where we pick only one representative among identical maximal sequences.

So, in summary, we aim at identifying a partition of the sequences in the alignment, where the sequences in each set of the partition have at least one maximum element (a sequence at least as informative as any other sequence in the set). One maximum from each set can be used as representatives to infer a ML tree, and then all the other elements of the sets can be attached to the corresponding maximum. A partition where the number of sets, and therefore the number of maxima, is equal to the number of maximal elements in the alignment (without counting duplicates), will be optimal, meaning that will have the minimum possible number of sets, and should therefore make ML inference the fastest. Note that, in this case, the set of maxima and the set of maximal elements (without duplicates) should coincide.

Note that from Lemma S1.1 above also follows the proof of the special case of the standard approach of removing only identical sequences.

#### S1.7 Estimating substitution rates

Substitution models are an essential component of ML phylogenetic inference, and we have described how we use a substitution rate matrix $Q$ to calculate phylogenetic likelihoods in the Main Text Methods. Here we describe how we estimate $Q$ during the estimation of the initial phylogeny. The same matrix $Q$ is then also used for the final topological improvements described below.

We have currently implemented three nucleotide substitution models in our software, the JC69[7], GTR[8] and UNREST[9] models. In the case of UNREST and GTR, we use as default initial values for the substitution rate matrix SARS-CoV-2 mutation rate estimates from[1], and we use as root nucleotide frequencies the nucleotide frequencies of the reference genome. In the case of GTR and UNREST, we update the substitution rates as more sequences are added to the tree (by default every 40 sequences). For each sequence we add, we record the number of substitutions of each type observed on the branch leading to the new sample; we only consider positions where the total likelihood genome list entry at the node of placement is one of the types $\mathbf{R}, \mathbf{A}, \mathbf{C}, \mathbf{G}$, and $\mathbf{T}$; where the same is the case for the partial likelihood vector of the new sample; and where these two types differ from each other. As an approximation, we ignore all other possible substitution events (for example those that might have happened as with a new placement an entry of type $\mathbf{O}$ at a tree node is connected to an entry of type **A** in the placed sample at a given site) which typically represent a minority of cases, and which are harder to label as substitutions. The total number of mutations of each type observed so far (plus initial pseudocounts) are used to update the substitution rates. For UNREST, mutation counts from **A** to **C** are used for the rate $q_{AC}$, and so on. For the reversible model GTR, mutation counts from **A** to **C** and from **C** to **A** are used for the rate $q_{AC}$, and so on. Each substitution rate is updated by setting it to the ratio of the new the number of mutations of the same type observed, over the root frequency (the frequency in the reference genome) of the source nucleotide of the considered substitution. Diagonal elements of the substitution rate matrix are defined so that rows of the matrix sum up to 0. The matrix is normalized so that the expected substitution rate is 1. The updated rates are then used to determine placement probabilities for new leaves. After the whole phylogenetic placement is concluded, and all samples have been added to the tree, all the genome lists are updated, and the substitution model is kept constant for the next inference stage comprising topological improvements (section S1.8).

#### S1.8 Tree topology improvement

We described so far how we build an initial tree by iteratively placing one sample at the time, and how we estimate a substitution rate matrix $Q$. In this section, we describe the second stage of the phylogenetic inference in MAPLE, where we improve the initial tree by modifying its topology and branch lengths.

Maximum likelihood phylogenetic inference methods usually employ topology-changing moves to a current tree in order to explore the phylogenetic tree space and attempt at finding tree topologies with highest likelihood. Typical tree topology change proposals are for example the nearest neighbour interchange ("NNI", a short-distance topology change proposal) scheme and subtree pruning and regrafting (SPR, a long-range topology change proposal) scheme[3]. Here, we propose an efficient implementation of the SPR scheme. The idea behind this approach is to re-adapt the methods for phylogenetic placement described in the Main Text Methods in order to re-place nodes of the phylogeny (either internal nodes or terminal ones) by severing them from the tree and proposing their placement elsewhere.

##### S1.8.1 Tree traversal loop for SPR initialization

To improve the initial tree, we traverse the tree from the root to the tips in preorder (first the parent node, then the left child and all its descendants, and finally the right child and all its descendants). At each non-root node $n$ we perform an SPR search procedure (described in detail in section S1.8.2) in which we sever the subtree rooted at $n$ from the current tree and attempt to regraft it somewhere else.

Before beginning the procedure, however, we evaluate the log-likelihood cost of the current placement of $n$ and its subtree — this corresponds to the total log-likelihood of the current tree, minus the log-likelihood of the subtree rooted at $n$ (but using improper root nucleotide frequencies all equal to 1.0) and the log-likelihood of its complement (the tree obtained by removing from the current tree the subtree rooted at $n$). This difference represents the contribution of the current placement of $n$ to the total likelihood of the current tree, and it can be calculated as described in section S1.3 without needing to calculate the individual components of the difference, and only using genome lists of node $n$ and its parent.

If this cost is below a certain threshold (default: $10^{-5}$ log-likelihood units), then the likelihood gain of a potential SPR move would be very limited and we do not initiate the SPR search procedure or any of the following steps. If the current cost is above the threshold, we first attempt at optimizing the length $l_n$ of the branch above $n$ at the current placement (branch lengths are coarsely optimized by recursively halving or doubling the initial length, as during placement). Given the best found value of $l_n$, this will define the current placement cost of $n$, and the SPR search will use branch length $l_n$ as a branch placement length for the other nodes of the tree as well. However, if the new current likelihood cost corresponding to $l_n$ is below our threshold, or if it is not but no SPR improvement is found, then the SPR procedure is converted into a simple branch length update move.

##### S1.8.2 SPR search

Likelihood costs for the current placement of a node $n$, and alternative placements, are calculated like the placement likelihood scores in section S1.3, with the difference that now the lower partial likelihoods that are being placed are not necessarily those of a sample, but are often those of an internal node.

Similarly to the placement search procedure, we traverse the tree in search of nodes and mid-branch points which would provide the best fit (highest likelihood) for a new placement/re-graft of $n$. However, the SPR search tree traversal is started at $n$, not at the root, and it does not traverse the subtree rooted at $n$. Another difference of our SPR search compared to our initial placement search is that now, when evaluating alternative placements, we have to consider the fact that $n$ needs to be first severed from the tree, which can affect the existing partial likelihood genome lists in the tree. For this reason, as we traverse the tree trying to re-place $n$, we also carry over a genome list representing new partial likelihoods at the considered node following the severing of $n$ from the tree. In other words, removing the subtree rooted at $n$ from the tree can affect ancestral nucleotide probabilities in the remainder of the tree, and we need to take this into account when searching a new attachment node for $n$ and its subtree.

For example, assume that $n$ and $n_2$ are child nodes of $P$, and so by severing $n$ from the tree, $P$ becomes a node with a single child node. Assume also that $P$ is the left child of its own parent node $P_2$, which we traverse first in order to assess it as a possible new placement of $n$. To do this, we cannot use the current total likelihood genome list of $P_2$, since it has been calculated considering also the data in the subtree rooted at $n$. Instead, we first calculate an alternative total likelihood genome list of $P_2$, following the potential severing of $n$, by combining the lower likelihood genome list of $n_2$ (which replaces the lower likelihood genome list of $P$) and the upper-right likelihoods genome list of $P_2$. We use this alternative total likelihood genome list to evaluate the placement cost of $n$ at $P_2$. We then also calculate an alternative lower likelihood genome list for $P_2$ by combining the lower likelihood of $n_2$ with the lower likelihood of the right child of $P_2$, and pass it on to the parent node of $P_2$ as we keep traversing the tree. Similarly, we calculate an alternative upper-left likelihood genome list for $P_2$ and pass it to its right child as we traverse it. At each step of tree traversal we therefore calculate some alternative genome lists, of which we use the total likelihood one for assessing the placement cost of $n$, while the others we pass on as we move further along the tree traversal. All these alternative genome lists do not immediately replace the old ones, but instead the old ones are stored in case no SPR improved re-grafting is found for $n$.

Usually, after a very few steps in this tree traversal, we find that the alternative partial likelihood genome lists coincide with the pre-existing ones (meaning that changes in the tree like the severing of a subtree typically only affect ancestral state probabilities for a small neighborhood near the severed node). When this happens, we avoid the calculation and passing on of alternative genome lists that would necessarily coincide with those already in the tree.

If, while traversing the tree during an SPR search, we reach a point at which the re-placement of $n$ is sufficiently unlikely (by default, more than 160 log-likelihood units more unlikely than the best placement found so far) and if the re-placement cost has increased by at least 1 log-likelihood unit a sufficient number of times (by default 4 times) while traversing the tree in the same direction, then we stop the SPR search in that direction.

Note that some of our heuristics for SPR search are similar to some that have been developed for other phylogenetic packages, and in particular RAxML. For example, some aspects of our SPR approach are similar to the Lazy Subtree Rearrangement[10] ("LSR"); however, we don't optimize three branch lengths at each SPR evaluation, but instead keep constant the sum of the lengths of two of the branches near the re-graft node; also, we don't define the SPR search radius based on the number of nodes traversed from the original location of the subtree, but instead we only use the difference of the log likelihood score of the proposed SPR moves against the original tree (approach that in itself is similar to the one by Stamatakis et al.[11]). This means that our SPR moves can potentially be more costly than LSR steps, since we could in principle explore a broader region of the tree, for example for subtrees containing extremely uninformative sequences, whose placement can be uncertain; at the same time, we usually avoid traversing areas of the tree where the re-placement of a subtree would be very unlikely.

##### S1.8.3 SPR move finalization

If the SPR search finds a better placement point than the current location of $n$, we first sever $n$ and its descendants from the tree and update all the genome lists in the rest of the tree accordingly (but again avoiding updating genome lists when the old one and the new are expected to coincide). Then, we place $n$ to its new location in the tree and we re-update the genome lists in the tree, starting from $n$ and traversing its subtree and the rest of the tree (but again stopping every time that new and old genome lists coincide). Thanks to the stopping criteria for these tree traversals, these genome list updates are usually not very demanding since only a small part of the tree is affected and therefore traversed.

While finalizing an SPR move, we mark as "dirty" each node whose genome lists have been modified. After the SPR search has been terminated for the whole tree, we re-traverse all the nodes that have been marked as dirty and attempt to re-place them again with SPR moves. This procedure is repeated until no genome lists are modified and therefore no node becomes dirty.

#### S1.9 Extended comparison with other phylogenetic inference methods

Sometimes specific phylogenetic method options used in can affect the computational performance and accuracy of phylogenetic inference. For this reason, to extend our comparison between phylogenetic methods, we considered a number of options for the different methods considered.

We test the performance of MAPLE using two settings:

- Default: it uses the default MAPLE options, which typically lead to a thorough exploration of tree space and accurate estimation.
- Fast: uses option "--fast" in MAPLE, which corresponds to setting 4 allowed failed moves per direction and log-likelihood threshold of 160 units during the initial placement, and to setting 2 allowed fails per direction and log-likelihood threshold of 80 during the SPR topological improvement stage. It also skips SPR search for nodes with a current likelihood cost below 1 log-likelihood unit.

The difference between the MAPLE default and fast options is that the default parameters lead to a more in-depth tree search, and so take longer but are also expected to result in trees with higher likelihoods.

For IQ-TREE 2 we considered three different speed settings:

- Fast (also considered in the main text): we used option "-fast", for which only nearest neighbour interchange (NNI) moves are used.
- Medium: we used default options and "-blmin 0.000000005" to allow shorter branch lengths.
- Slow: in addition to "-blmin 0.000000005" we also used options "-nstop 500" to increase the number of unsuccessful iterations before stopping, and "-pers 0.1" to decrease the default perturbation strength.

For simulated data with rate variation we used in IQ-TREE 2 a GTR+G substitution model instead of a GTR one.

For FastTree 2 we used four speed settings:

- Fastest (also considered in the main text): we used option "-fastest" to reduce the time demand of NNI steps.
- Fast: default setting.
- Medium: we used default options and "-spr 4" to increase the number of rounds of minimum-evolution SPR moves.
- Slow: in addition to "-spr 4" we also used options "-mlacc 2 -slownni" to make the maximum-likelihood NNIs more exhaustive.

For simulations with rate variation we used option "-cat 4" instead of "-nocat".

RAxML-NG we considered two speed settings:

- Fast (also considered in the main text): we used option "--blmin 0.000005" to increase the minimum branch length considered and option "--tree pars1" to start the tree search from a parsimony tree.

- Slow: we used option "- -blmin 0.000000005 - -tree pars3" to decrease the minimum branch length and start the tree search from 3 parsimony trees.

For simulations with rate variation we use a GTR+G model instead of GTR.

In addition to RAxML-NG, we also ran RAxML v8.2.11 (raxmlHPC)[12] using options "-F" to skip gamma model estimation, and "-c 1 -V" to skip the model of rate variation. We ran RAxML with two speed settings:

- Fast: we used option "-D" to speed up tree search.
- Slow: default settings.

For simulations with rate variation we used options "-c 4 -m GTRCAT".

matOptimize and UShER were run either one after the other (as considered in the main text), or, as a faster and less memory intensive alternative, we ran UShER not followed by matOptimize topological improvements.

### Supplementary Methods References


1. De Maio, N. *et al.* Mutation rates and selection on synonymous mutations in SARS-CoV-2. *Genome Biology and Evolution* **13,** evab087 (2021).
2. Boussau, B. & Gouy, M. Efficient likelihood computations with nonreversible models of evolution. *Systematic Biology* **55,** 756–768 (2006).
3. Swofford, D., Olsen, G., Waddell, P. & Hillis, D. *Phylogeny Inference* 407–514 (Sinauer Associates, Massachusetts, 1996).
4. Yang, Z., Kumar, S. & Nei, M. A new method of inference of ancestral nucleotide and amino acid sequences. *Genetics* **141,** 1641–1650 (1995).
5. Minh, B. Q., Lanfear, R., Trifinopoulos, J., Schrempf, D. & Schmidt, H. IQ-TREE version 2.1. 2: Tutorials and Manual Phylogenomic software by maximum likelihood (2021).
6. Felsenstein, J. Confidence limits on phylogenies: an approach using the bootstrap. *Evolution* **39,** 783–791 (1985).
7. Jukes, T. H., Cantor, C. R., *et al.* Evolution of protein molecules. *Mammalian Protein Metabolism* **3,** 21–132 (1969).
8. Tavaré, S. Some probabilistic and statistical problems in the analysis of DNA sequences. *Lectures on Mathematics in the Life Sciences* **17,** 57–86 (1986).
9. Yang, Z. Estimating the pattern of nucleotide substitution. *Journal of Molecular Evolution* **39,** 105–111 (1994).
10. Stamatakis, A., Ludwig, T. & Meier, H. RAxML-III: a fast program for maximum likelihood-based inference of large phylogenetic trees. *Bioinformatics* **21,** 456–463 (2005).
11. Stamatakis, A., Blagojevic, F., Nikolopoulos, D. S. & Antonopoulos, C. D. Exploring new search algorithms and hardware for phylogenetics: RAxML meets the IBM cell. *The Journal of VLSI Signal Processing Systems for Signal, Image, and Video Technology* **48,** 271–286 (2007).
12. Stamatakis, A. RAxML version 8: a tool for phylogenetic analysis and post-analysis of large phylogenies. *Bioinformatics* **30,** 1312–1313 (2014).
